# Recurrent hypoxia in a rat model of sleep apnea during pregnancy leads to microglia-dependent respiratory deficits and persistent neuroinflammation in adult male offspring

**DOI:** 10.1101/2022.12.20.521336

**Authors:** Carly R. Mickelson, Andrea C. Ewald, Maia G. Gumnit, Armand L. Meza, Abigail B. Radcliff, Stephen M. Johnson, Jonathan N. Ouellette, Bailey A. Kermath, Avtar S. Roopra, Michael E. Cahill, Jyoti J. Watters, Tracy L. Baker

**Author notes:** co-corresponding Correspondence to: Tracy L. Baker, PhD, 2015 Linden Dr., Madison, WI 53706, Jyoti J. Watters, PhD, 2015 Linden Dr., Madison, WI 53706. contributed equally (Carly R. Mickelson, Andrea C. Ewald, Maia G. Gumnit, Armand L. Meza).

## Abstract

Sleep apnea (SA) during pregnancy is detrimental to the health of the pregnancy and neonate, but little is known regarding long-lasting consequences of maternal SA during pregnancy on adult offspring. SA is characterized by repeated cessations in breathing during sleep, resulting in intermittent hypoxia (IH). We show that gestational IH (GIH) in rats reprograms the male fetal neuroimmune system toward enhanced inflammation in a region- and sex-specific manner, which persists into adulthood. Male GIH offspring also had deficits in the neural control of breathing, specifically in the ability to mount compensatory responses to central apnea, an effect that was rescued by a localized anti-inflammatory or microglial depletion. Female GIH offspring appeared unaffected. These results indicate that SA during pregnancy sex- and region-dependently skews offspring microglia toward a pro-inflammatory phenotype, which leads to long-lasting deficits in the capacity to elicit important forms of respiratory neuroplasticity in response to breathing instability. These studies contribute to the growing body of recent evidence indicating that SA during pregnancy may lead to sex-specific neurological deficits in offspring that persist into adulthood.

## INTRODUCTION

It is both scientifically thrilling and practically terrifying that a mother’s experiences during pregnancy can contribute to disease development in her adult offspring (Bilbo et al., 2018). For example, prolonged or severe hypoxia during perinatal life is associated with an increased risk for neurological disorders in offspring (Amgalan et al., 2021), such as autism spectrum disorder (ASD), schizophrenia or intellectual disability (Dalman et al., 2001; Modabbernia et al., 2016); typically this form of hypoxia is associated with a traumatic or disordered pregnancy and physicians are well-aware of risks to the child. Underappreciated is that many women commonly experience recurrent episodes of brief hypoxia during pregnancy in the form of sleep apnea (SA). SA is characterized by repeated pauses in breathing, and occurs in 10-30% of all pregnancies by the third trimester (Pien et al., 2014; Lockhart et al., 2015). Although SA in pregnancy is associated with adverse consequences to the neonate, including preterm birth and NICU admission (Ding et al., 2014), little is known regarding long-lasting effects of maternal SA on adult offspring. Recent evidence in animal models indicates that intermittent hypoxia (IH) associated with SA leads to life-long alterations in offspring physiology, with male offspring more severely affected. Adult male offspring born from rodent dams exposed to IH exhibit deficits in metabolism (Khalyfa et al., 2017; Cortese et al., 2021), cardiovascular function (Song et al., 2021) and have behavioral deficits characteristic of ASD in humans (Vanderplow et al., 2022). However, effects on other core physiological systems are unknown.

Sex-specific reprogramming of the neuroimmune response toward a persistently pro-inflammatory state is a common outcome of an adverse *in utero* environment and may contribute to offspring cognitive deficits (Al-Haddad et al., 2019). Neuroinflammation in brain regions important in cognition undermines synaptic plasticity (Rizzo et al., 2018), a foundational piece of cognitive processing that enables the CNS to adapt to new experiences over the lifetime. Microglia play critical roles in creating and resolving neuroinflammation in the healthy and injured brain (Bilbo et al., 2018; Oh et al., 2020; Yegla et al., 2021). While the detrimental impact of an adverse perinatal environment on offspring cognitive function has been intensely investigated, potential effects of the perinatal environment on CNS regions important in breathing remain poorly understood.

Breathing is a complex neuromotor behavior that relies on a precisely coordinated pattern of muscle activation to hold the airway open and expand the thoracic cavity. Plasticity is a key element of a healthy respiratory control system since it adjusts muscle activation patterns to optimize breathing throughout the lifetime (Fuller and Mitchell, 2017). Neural circuits driving breathing are tuned for near constant activity from birth until death, and as such, are exquisitely sensitive to reductions in respiratory neural activity, even in the absence of a change in blood gases. Indeed, central hypopnea/apnea trigger a chemoreflex-independent, proportional enhancement in inspiratory motor output to muscles maintaining upper airway tone and expanding the thoracic cavity, a form of plasticity known as inactivity-induced inspiratory motor facilitation (iMF) (Braegelmann et al., 2017). Although respiratory neural activity is likely monitored in multiple neural circuits, local mechanisms operating within inspiratory motor neuron pools are key for iMF initiation (Streeter and Baker-Herman, 2014a).

We modeled SA during pregnancy by exposing rat dams to IH during their sleep phase in late gestation (gestational intermittent hypoxia; GIH). We report that GIH leads to long-lasting respiratory control and neuroimmune dysregulation in adult male, but not female, offspring. Adult male GIH offspring exhibit enhanced microglial inflammatory gene expression in spinal regions encompassing the phrenic motor pool, which interferes with the capacity to trigger iMF in response to recurrent central apnea. GIH-induced neuroinflammation was CNS-region specific, with no evidence for brainstem neuroinflammation. The capacity to elicit iMF could be rescued by a spinal anti-inflammatory or microglial depletion. Our results indicate that GIH sex- and region-specifically skews microglia toward a pro-inflammatory phenotype, impairing important forms of respiratory neuroplasticity elicited by breathing instability.

## RESULTS

### GIH impairs respiratory neuroplasticity

We first tested the hypothesis that the capacity to elicit respiratory neuroplasticity is impaired in adult offspring exposed to GIH compared to gestational intermittent normoxia (GNX)-exposed controls (Experimental Series 1). To induce respiratory plasticity, inspiratory neural activity was briefly silenced five times for ∼1min each to mimic recurrent central apnea; rats continued to be ventilated during the central apnea so that no hypoxia or hypercapnia was experienced. Representative compressed phrenic neurograms show phrenic inspiratory motor output at baseline, during, and for 60 min following recurrent central apnea (**Fig. 1a**). Phrenic inspiratory activity was monitored for an equivalent duration in “time controls” exposed to the same surgical preparation, but with no central apneas. Recurrent central apneas triggered a compensatory increase in phrenic inspiratory burst amplitude in both male (59.3±4.8% baseline, n=11, p<0.0001) and female (78±12.9% baseline, n=10, p=0.0036) GNX offspring relative to time controls (13.3±5.1% baseline, n=13), indicating iMF (**Fig. 1a, 1b**). Likewise, recurrent central apnea triggered robust iMF in female GIH offspring (63.6±11.7% baseline, n=9, p=0.0150) that was not statistically different from female GNX offspring (p=0.9688; **Fig. 1b**). Strikingly, recurrent central apnea did not elicit iMF in male GIH offspring (3.6±6.8% baseline, n=9, p=0.8627 compared to time controls; p<0.0001 compared to male GNX offspring). These data show that normal compensatory responses to reductions in respiratory neural activity are impaired by GIH in male, but not female, offspring.

**Figure 1:**
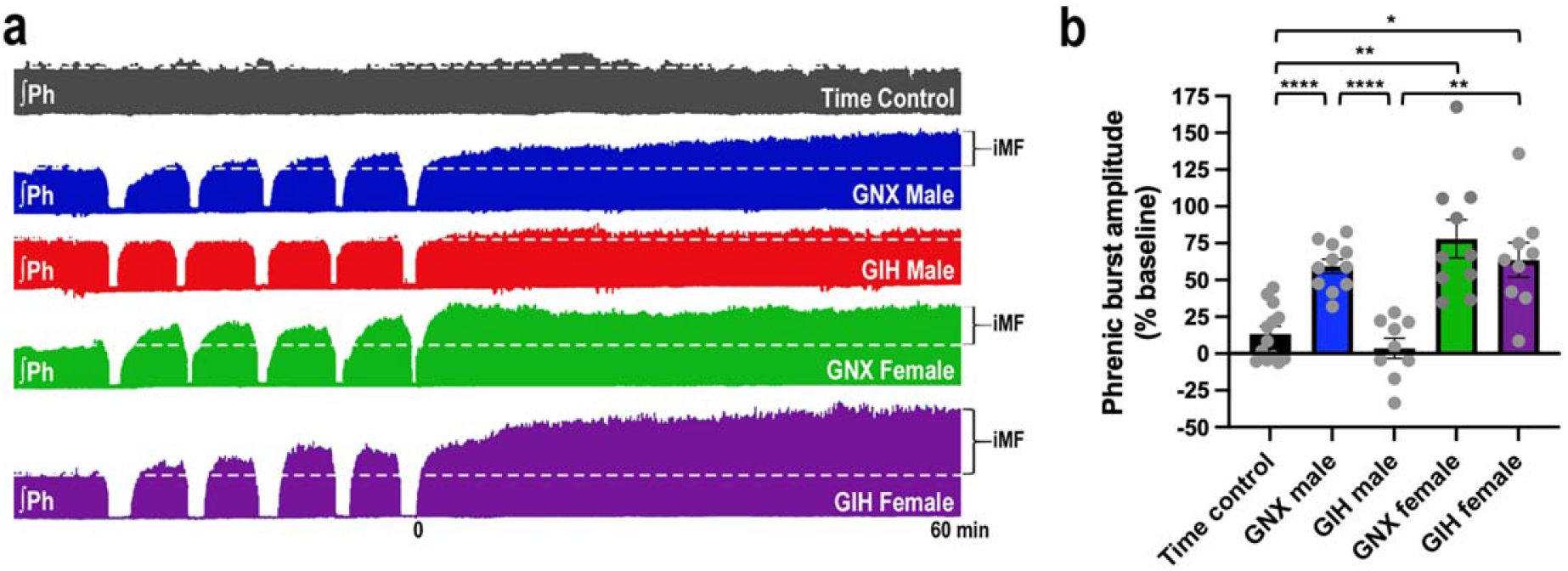
Gestational intermittent hypoxia (GIH) dysregulates compensatory respiratory neuroplasticity in adult male offspring. **(a)** Representative compressed phrenic neurograms depicting phrenic burst amplitude before, during, and for 60 minutes following exposure to recurrent reductions in respiratory neural activity (5, ∼1 min central apneas) or the equivalent duration in a rat not receiving central apnea (time control). Dotted white line represents baseline amplitude. **(b)** Average percent change (±SEM) in the amplitude of phrenic inspiratory output from baseline at 60 minutes following the fifth apnea. Treatment groups exposed to central apnea were compared to time controls that did not receive central apnea (n=13) to determine if respiratory neural activity deprivation triggered plasticity. In adult male and female rats exposed to gestational intermittent normoxia (GNX; males n=11, t=6.535, df=21.98, p<0.0001; females n=10, t=4.664, df=11.85; p=0.0036) and female GIH rats (n=9, t=3.929, df=11.08, p=0.0150), recurrent reductions in respiratory neural activity elicited a compensatory increase in phrenic inspiratory burst amplitude, indicating iMF. There was no statistical difference between the magnitude of iMF in GNX versus GIH females (t=0.8257, df=16.98, p<0.9688). Conversely, recurrent central apnea did not elicit increased phrenic inspiratory burst amplitude in adult male GIH rats (n=9, t=1.135, df=16.12, p=0.8627 relative to time controls; t=4.664, df=14.99, p<0.0001 relative to male GNX offspring), indicating that the ability to trigger compensatory enhancements in phrenic inspiratory output in response to respiratory neural activity deprivation (i.e., iMF) is impaired in adult male, but not female, offspring of dams exposed to intermittent hypoxia during gestation. Statistical analysis: Welch ANOVA (W_(4,21.64)_ = 18.63, p<0.0001) followed by Dunnett’s T3 multiple comparisons test, *p<0.05, **p<0.005, ****p<0.0001).

### GIH does not impact maternal care

Because other models of prenatal insult show changes in maternal care (Moore and Power, 1986), which can impact respiratory control in the offspring (Genest et al., 2007), we tested the hypothesis that GIH impacted maternal care. Several aspects of rodent maternal care were measured, including the time it took GNX and GIH mothers to approach their pups after they had been removed for 10 mins (GNX n=6, GIH n=6; **Suppl. Fig. 1a)**, retrieve the first pup (GNX n=11, GIH n =10; **Suppl. Fig. 1b)**, retrieve all pups (GNX n=11, GIH n=10; **Suppl. Fig. 1c)**, begin licking, grooming or sniffing the pups (GNX n=6, GIH n=6; **Suppl. Fig. 1d)** and begin crouching after all pups had been retrieved (GNX n=6, GIH n=5; **Suppl. Fig. 1e)**. We detected no differences in these measures of maternal care between GIH and GNX dams, suggesting that poor maternal care does not likely play a role in GIH-induced deficits in adult male offspring.

### GIH increases spinal cord, but not brainstem, inflammation in male offspring

To test whether GIH induces neuroinflammation in CNS regions underlying breathing, we measured inflammatory gene expression in tissue homogenates from brainstem, where respiratory rhythm is generated, and cervical spinal cord segments C3-C6, where phrenic motor neurons reside. We focused on several general inflammatory markers that are increased in humans and rodent models with deficits in respiratory control, including interleukin-1 beta (IL-1β), s100a8 (a calcium binding protein), C-X-C motif chemokine ligand 10 (CXCL10), Janus kinase 1 (JAK1), and signal transducer and activator of transcription 1 (STAT1) (Jain et al., 2012; Anderson et al., 2014; Almatroodi et al., 2015). For male GIH offspring, no differences in brainstem inflammatory markers were detected **(Fig. 2a)** but increases in *Il1β* (GNX n=8, GIH n=7, p=0.04), *s100a8* (GNX n=7, GIH n=6, p=0.019), *Cxcl10* (GNX n=6, GIH n=6, p=0.078), *Jak1*, (GNX n=7, GIH n=5, p=0.046) and *Stat*1 (GNX n=6, GIH n=6, p=0.065) gene expression were observed in cervical spinal cord **(Fig. 2b)**. In contrast, *Il1β* and *Cxcl10* gene expression was reduced in adult female GIH offspring in both brainstem (*Il1β* GNX n=8, GIH n=7, p=0.035; *s100a8* GNX n=7, GIH n=6, p=0.11; *Cxcl10* GNX n=7, GIH n=7, p=0.02) **(Fig. 2c)** and cervical cord (*Il1β* GNX n=8, GIH n=7, p=0.019; *s100a8* GNX n=7, GIH n=8, p=0.17; *Cxcl10* GNX n=6, GIH n=7, p=0.04), with a trend toward a reduction in *s100a8* (brainstem: GNX n=7, GIH n=6, p=0.11; cervical cord: GNX n=7, GIH n=8, p=0.17) **(Fig. 2d)**. These observations underscore critical sex differences in how offspring adapt to the maternal IH insult and indicate that spinal inflammation may play a role in GIH-induced neuroplasticity impairments.

**Figure 2:**
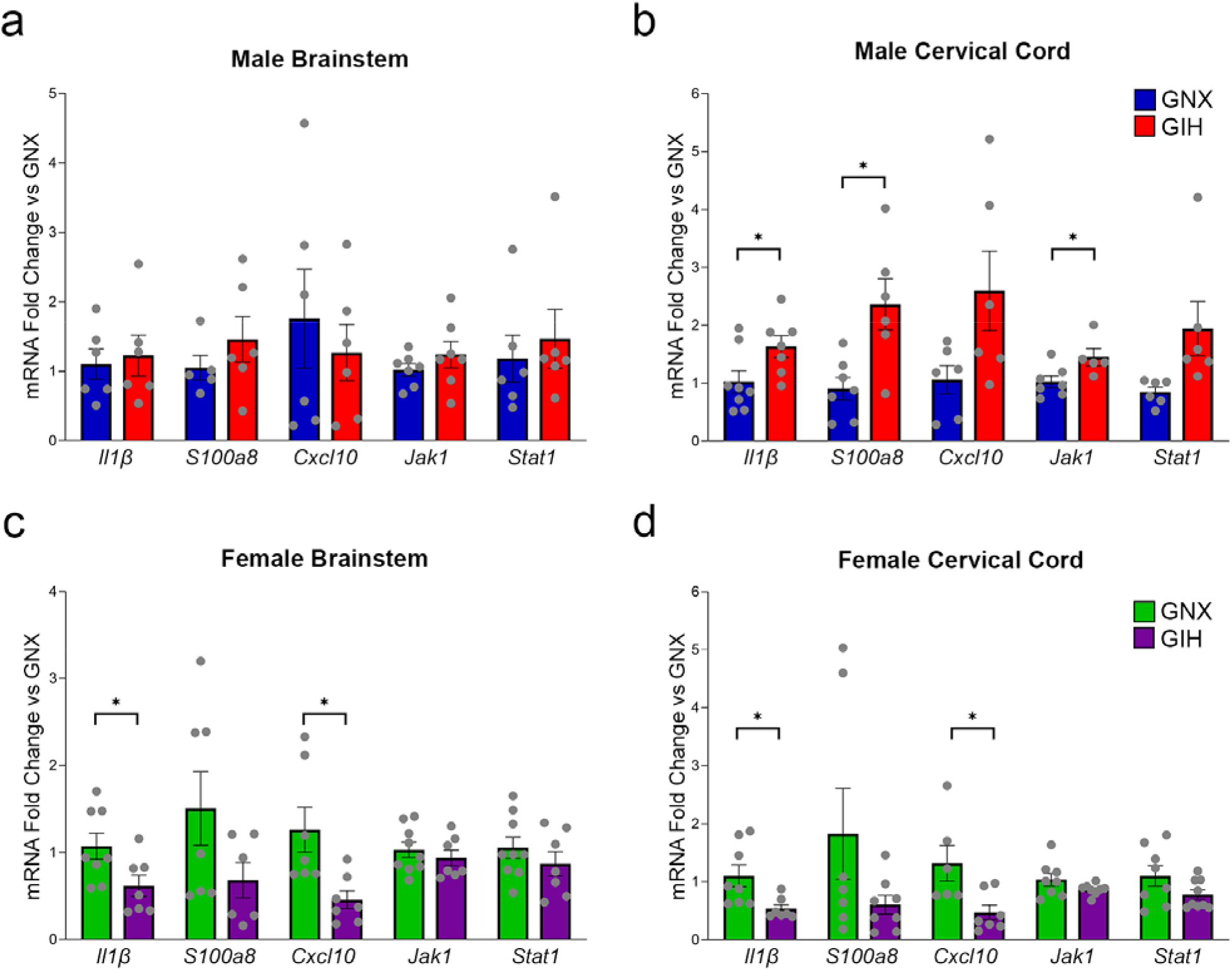
GIH males have increased inflammation in cervical spinal cord. **(a, b)** Basal inflammatory gene expression was analyzed in adult male brainstem **(a)** and cervical spinal cord **(b)** tissue homogenates using an unpaired t-test with Welch’s correction. Increases in *Il1β* (GNX,GIH n=8,7; t=2.285, df=12.94, F=1.170, p=0.04), *s100a8* (GNX,GIH n=7,6; t=3.029, df=6.885, F=4.482, p=0.019), *Cxcl10* (GNX,GIH n=6,6; t=2.101, df=6.249, F=7.876, p=0.078), *Jak1*, (GNX,GIH n=7,5; t=2.395, df=7.391, F=1.598, p=0.046) and *Stat*1 (GNX,GIH n=6,6; t=2.315, df=5.351, F=28.45, p=0.065) gene expression were seen in male spinal cord but not in brainstem. **(c, d)** Basal inflammatory gene expression was analyzed in adult female brainstem **(c)** and cervical spinal cord **(d)** tissue homogenates using an unpaired t-test with Welch’s correction. In contrast to male offspring, decreases in *IL-1β, S100a8, Cxcl10* gene expression were seen in both female brainstem (*Il1β* GNX,GIH n=8,7; t=2.361, df=12.81, F=1.706, p=0.035; *s100a8* GNX,GIH n=7,6; t=1.763, df=8.503, F=5.149, p=0.11; *Cxcl10* GNX,GIH n=7,7; t=2.905, df=7.898, F=6.161, p=0.02) and cervical spinal cord (*Il1β* GNX,GIH n=8,7; t= 2.852, df=8.822, F=8.505, p=0.019; *s100a8* GNX,GIH n=7,8; t=1.537, df=6.5, F=21.02, p=0.17; *Cxcl10* GNX,GIH n=6,7; t=2.538, df=6.702, F=4.987, p=0.04). Collectively, these data demonstrate sex- and CNS region-specific changes in inflammatory gene expression induced by GIH exposure. *p<0.05 relative to GNX controls.

### Adult offspring brainstem and spinal microglial transcriptomes are differentially reprogrammed by GIH

Since microglia are important mediators of neuroinflammation, we next examined whether GIH skewed microglia toward a pro-inflammatory phenotype. RNA-sequencing was performed on microglia (CD11b+ cells) immunomagnetically isolated from brainstem and cervical spinal cord (C3-C6; n=5/treatment). Principal component analysis of brainstem and cervical spinal microglia transcriptomes revealed little variability between male GNX and GIH treatments in either region **(Fig. 3a)**, although there were notable differences between brainstem and cervical spinal cord transcriptomes. Surprisingly, in cervical spinal microglia from female GIH offspring, there were nearly 4000 differentially expressed genes, with only modest changes in brainstem microglial genes **(Suppl. Fig 2**). Our subsequent analyses focused on transcriptomic changes in male GIH offspring microglia as our interest was in deficits in respiratory plasticity. However, understanding the physiological role for microglial transcriptomic changes in GIH females represents an exciting direction for future study. In isolated brainstem microglia from male GIH offspring, only 6 out of the 13,504 expressed genes were differentially expressed (4 up, 2 down) **(Fig. 3b; Table 1)**. Similarly, of the 12,982 genes expressed in male GIH offspring cervical spinal microglia, only 29 were differentially expressed (6 up, 23 down) (FDR < 0.05; **Fig 3c; Table 1**). Despite the relatively low number of differentially expressed genes, all of the known differentially upregulated genes in the cervical spinal cord are putative targets of either NF-κB or STAT transcription factors (**Fig. 3d**). Notably, the IκB kinase epsilon isoform (Iκbkε) was 1 of the 6 upregulated transcripts in spinal microglia from male GIH offspring. Iκbkε targets the NF-κB inhibitory subunit (IκB) for degradation by the proteasome, resulting in activation of NF-κB-dependent inflammatory signaling, consistent with the enhanced neuroinflammatory gene expression data we observed in Figure 2.

**Table 1:**
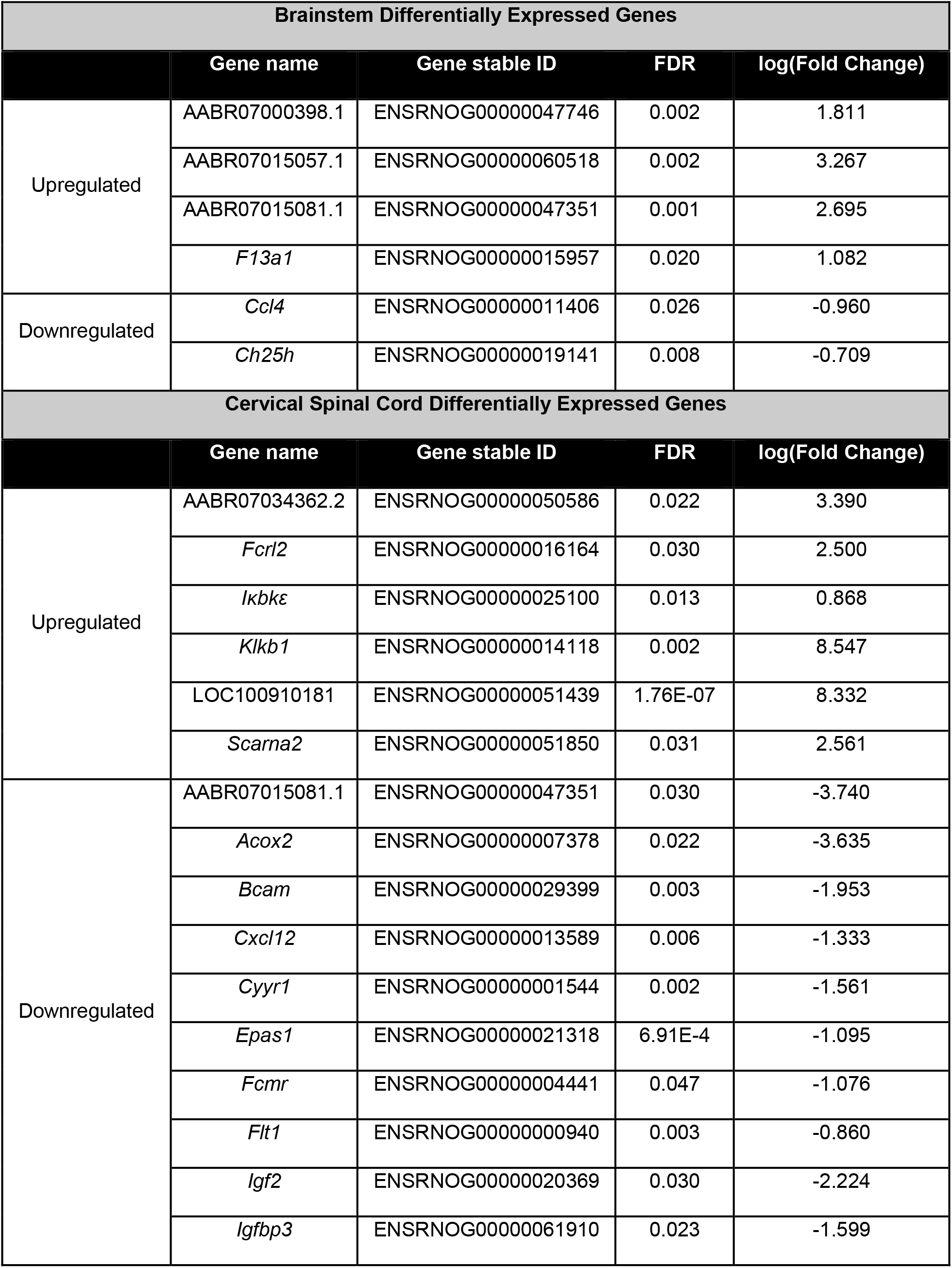
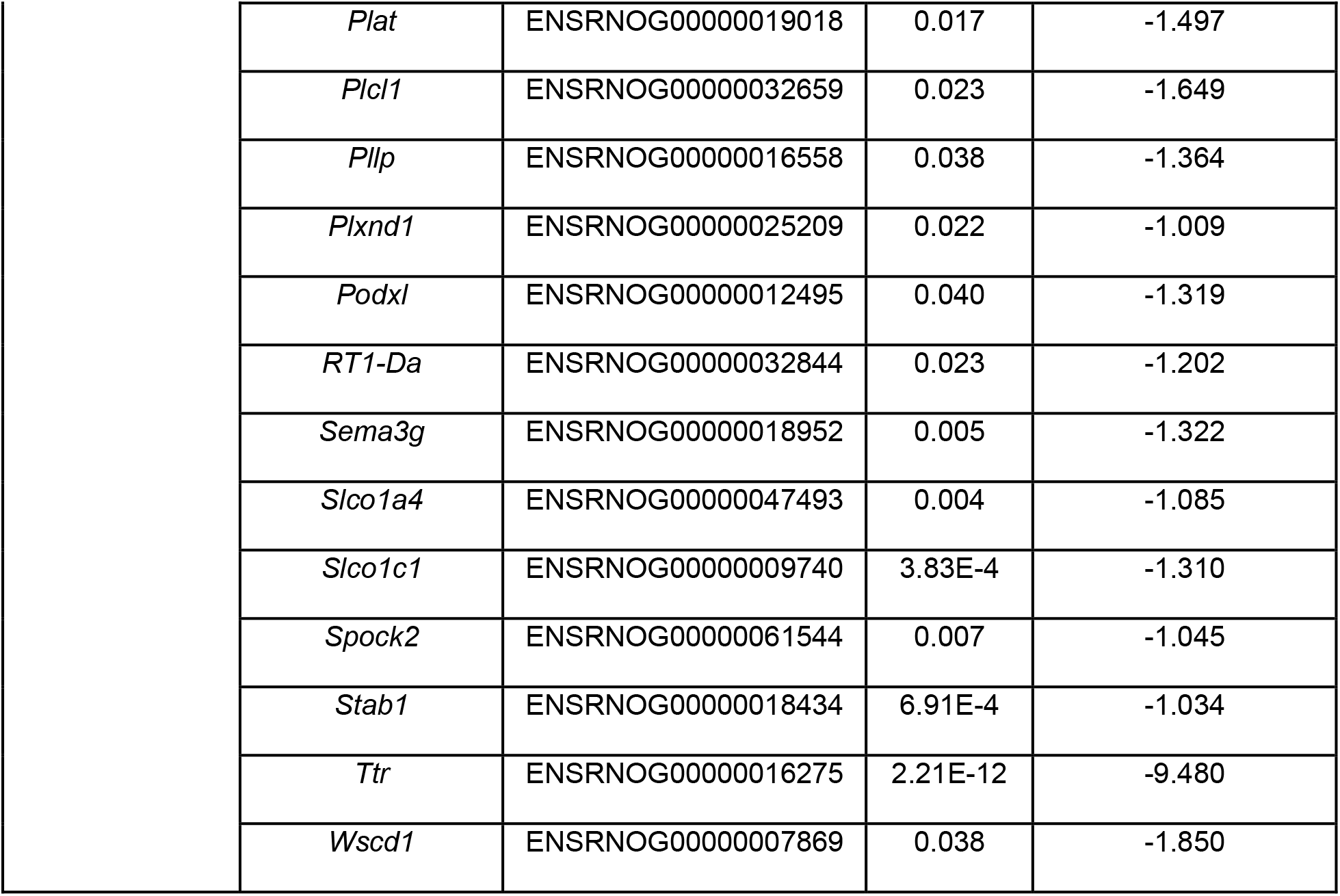
Differentially Expressed Genes.

**Figure 3:**
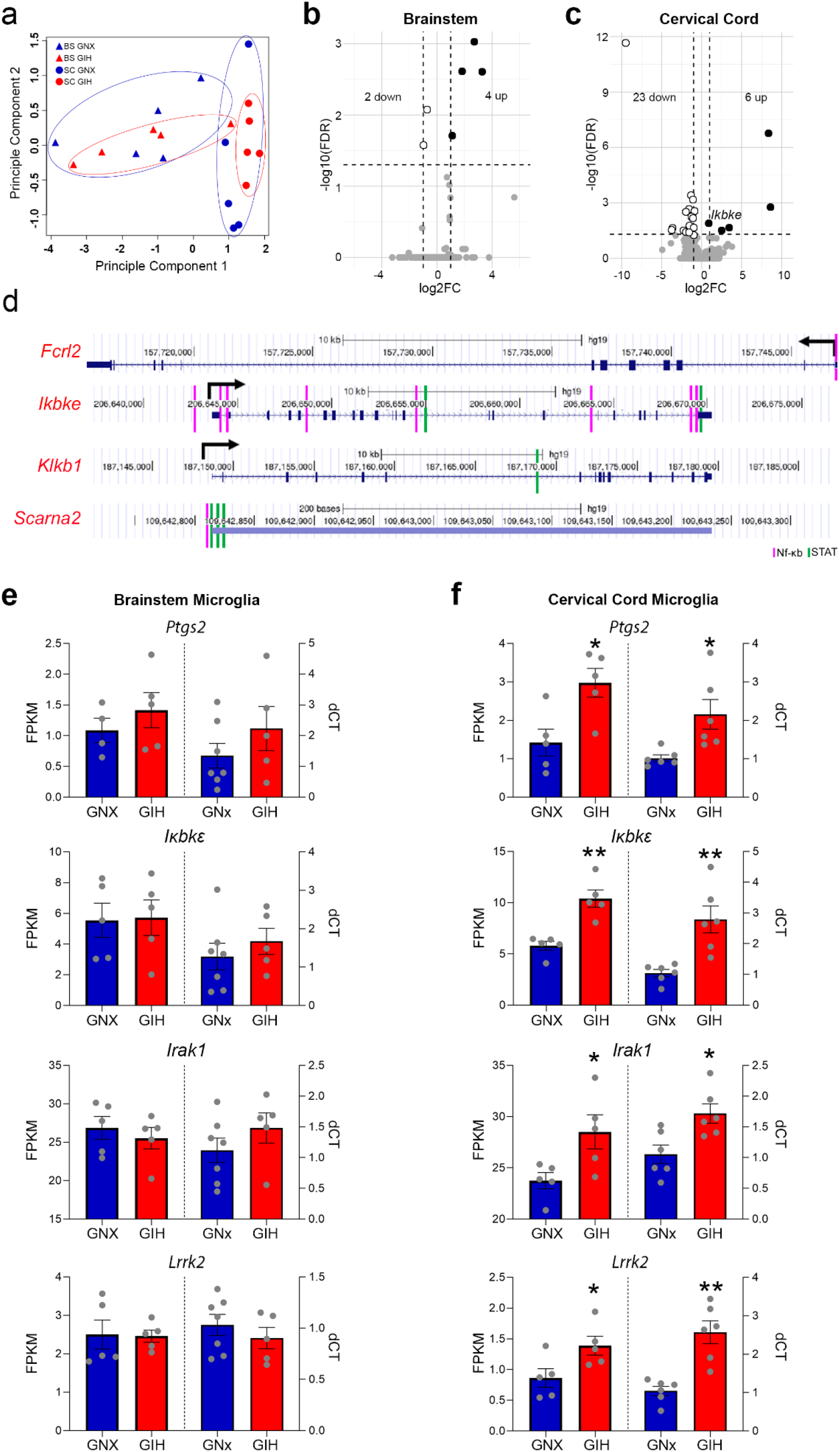
GIH differentially upregulates inflammatory gene transcripts in male spinal microglia. **(a)** Principal component analysis of RNA-seq data from male brainstem and cervical spinal microglia (n = 5/treatment) show greater transcriptomic differences resulting from CNS region than by GIH treatment. **(b)** Volcano plot indicating differential gene expression in microglia isolated from the male GIH offspring brainstem. Few genes (6) were altered by GIH. Black dots represent significantly upregulated genes (FDR < 0.05); white dots represent significantly downregulated genes. FC, fold change. **(c)** Volcano plot of differential gene expression in male GIH offspring cervical spinal microglia. 29 genes were differentially expressed; black dots represent significantly upregulated genes (FDR < 0.05); white dots represent significantly downregulated genes. **(d)** Gene tracks from the ENCODE database of the 4 known upregulated genes in GIH male spinal microglia demonstrate binding sites for NF-κB (pink) and STAT (green) below each track, suggesting that differentially upregulated gene expression in GIH spinal microglia may be regulated by NF-κB and STAT transcription factor activity. **(e**,**f)** To ascertain evidence of enhanced basal inflammation in GIH offspring microglia, FPKM (fragments per kilobase of exon per million mapped fragments) analyses of RNA-seq data (left) and RT-qPCR (right) for the inflammatory genes Ptgs2, Ikbke, Irak1, and Lrrk2 were performed. **(e)** No differences in the expression of Ptgs2, Ikbke, Irak1, or Lrrk2 were observed in male brainstem microglia but **(f)** enhanced inflammatory gene expression for Ptgs2, Ikbke, Irak1 and Lrrk2 was observed in male GIH spinal microglia (Ptgs2 FPKM t=3.029, df=7.96, F=1.152, p=0.016 and dCT t=2.888, df=5.489, F=20.40, p=0.031; Ikbke FPKM t=4.836, df=5.986, F=3.762, p=0.003 and dCT t=3.867, df=5.777, F=12.80, p=0.009; Irak1 FPKM t=2.581, df=5.690, F=4.512, p=0.044 and dCT t=3.04, df=9.985, F=1.08, p=0.013; Lrrk2 FPKM t=2.439, df=7.998, F=1.034, p=0.041 and dCT t=4.766, df=6.636, F=5.945, p=0.002). *p<0.05, **p<0.01 relative to GNX controls.

We evaluated the FPKM values of all expressed genes within the male brainstem and cervical cord microglia RNA-Seq datasets. Interrogating for genes specifically associated with the Gene Ontology search terms “NF-κB signaling” (GO: 0038061) and “neuroinflammatory response” (GO: 0150076) we found the NF-κB target genes prostaglandin-endoperoxide synthase 2 (Ptgs2, or COX-2), Iκbkε, interleukin-1-receptor-associated-kinase-1 (Irak1), and leucine-rich repeat kinase 2 (Lrrk2) to be increased. In brainstem microglia, there were no detectable changes induced by GIH in the expression of any of these genes either by RNA-Seq FPKM **(Fig. 3e, left)** or qPCR analyses **(Fig. 3e, right)**. However, in cervical spinal microglia, GIH increased the expression of all four of these inflammatory genes both by RNA-Seq FPKM **(Fig. 3f, left)** and individual qPCR confirmation analyses **(Fig. 3f, right)**, indicating that NF-κB signaling pathways may be hyperactivated in male GIH cervical spinal microglia.

### GIH-induced spinal neuroinflammation impairs iMF in adult male GIH offspring

Since iMF is a predominantly spinal form of neuroplasticity (Streeter and Baker-Herman, 2014a) and we observed increased neuroinflammation specifically in the spinal cord, we tested whether functional inhibition of local spinal inflammation, and NF-κB/STAT signaling in particular, could rescue iMF expression in adult male GIH offspring (Experimental Series 2). We intrathecally delivered vehicle (DMSO) or TPCA-1 (potent inhibitor of Iκ-kinases and STAT transcription factor activation) directly over the cervical spinal segments C3-C6, where phrenic motor neurons reside, prior to administering recurrent central apneas (**Fig. 4a, b**). Intrathecal TPCA-1 did not alter iMF magnitude in male GNX offspring (vehicle: 60.3±10.4% baseline, n=9; TPCA-1: 59.7±10.2% baseline, n=6; p>0.9999). As in figure 1, iMF was significantly impaired in vehicle-treated adult male GIH offspring (26.5±3.1%, n=8) relative to vehicle-treated GNX offspring (p=0.048). Interestingly, local spinal TPCA-1 treatment in male GIH offspring restored the capacity to express iMF in response to recurrent central apnea (75.6±10.8%, n=7, p=0.004 relative to vehicle GIH) to a level similar to that observed in GNX offspring (p=0.6726). These data indicate that GIH-induced spinal neuroinflammation abolishes the ability to elicit compensatory increases in phrenic inspiratory output in response to reductions in respiratory neural activity in adult male GIH offspring.

**Figure 4:**
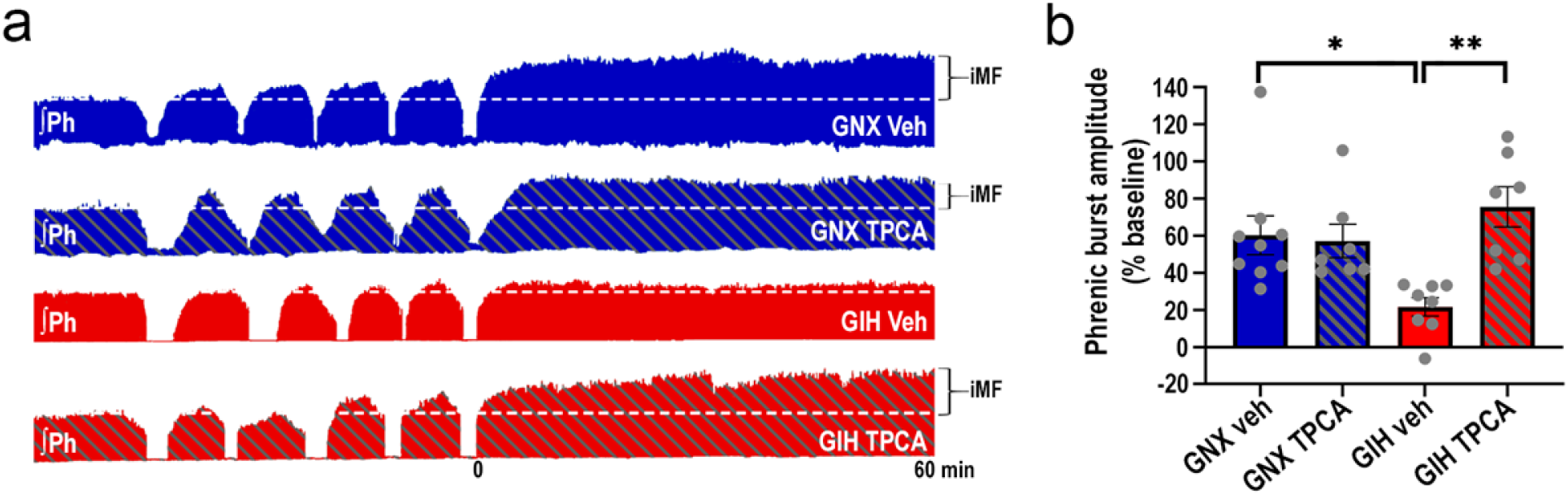
TPCA-1 administration to the phrenic motor pool restores iMF. **(a)** Representative compressed phrenic neurograms depicting phrenic burst amplitude before, during, and for 60 minutes following exposure to recurrent reductions in respiratory neural activity (5, ∼1 min neural apneas). Dotted white line represents baseline amplitude. ∼45 minutes prior to respiratory neural activity deprivation, rats received intrathecal injections of the Iκ-kinases/STAT inhibitor TPCA-1 or intrathecal vehicle in spinal regions encompassing the phrenic motor pool. **(b)** Average percent change (±SEM) in phrenic inspiratory burst amplitude from baseline at 60 minutes following the fifth central apnea. 2-way ANOVA revealed a significant effect of TPCA treatment (F_(1,26)_ =6.82, p=0.0148) and a significant interaction between TPCA treatment and GIH/GNX status (F_(1,26)_=7.132, p=0.0129). Tukey’s post-hoc test revealed that there was no difference in the magnitude of iMF between GNX rats treated with vehicle or TPCA-1 (n=9, 6 respectively; p>0.9999). Similar to previous results, iMF magnitude in vehicle treated male GIH rats (n=8) was significantly lower than the response in vehicle treated GNX males (p=0.0480). However, local application of TPCA-1 to the phrenic motor pool of male GIH (n=7) offspring rescued the capacity to trigger compensatory increases in phrenic inspiratory output following recurrent central apneas (p=0.004 relative to vehicle treated GIH; p=0.6726 relative to TPCA treated GNX). Collectively, these data demonstrate that GIH-induced spinal inflammation impairs the ability to elicit compensatory increases in phrenic inspiratory output in response to respiratory neural activity deprivation in adult male GIH offspring. *p<0.05, **p<0.01.

### Aberrant microglial activities play a key role in impairing iMF in adult male GIH offspring

To determine whether aberrant microglial function, specifically, contributes to neuroinflammation-induced impairments in iMF, rats were treated with vehicle or the CSF1R inhibitor Pexidartinib (PLX3397; 80mg/kg daily for seven days, p.o.) to pharmacologically deplete microglia **(Fig. 5a-o)**. Microglial depletion was confirmed using immunohistochemistry. Compared to vehicle-treated control groups, PLX3397 significantly reduced Iba+ cell numbers by ∼73% in the ventral horn (near phrenic motor neurons) of both GNX (Vehicle n=4, PLX n=6, p<0.0001) and GIH (Vehicle n=4, PLX n=5, p<0.0001) offspring **(Fig. 5b, d, m)**. Similar reductions were observed in the brainstem (not shown). PLX administration affected neither GFAP+ area fluorescence in GNX (Vehicle n=3, PLX n=5, p=0.424) or GIH offspring (Vehicle n=4, PLX n=3, p=0.682) **(Fig. 5f, h, n)** nor NeuN+ cell counts in GNX (Vehicle n=3, PLX n=6, p=0.754) or GIH offspring (Vehicle n=4, PLX n=5, p=0.895) **(Fig. 5j, l, o)**. Consistent with other recent reports in the rat brain (Oh et al., 2020; Yegla et al., 2021), these results confirm that seven days of PLX3397 treatment significantly decreased microglial numbers in the rat cervical spinal cord, without detectably impacting astrocyte activation status and neuron populations. To assess for evidence of astrogliosis in the brainstem, a key site involved in the neural control of breathing, we quantified GFAP+ area fluorescence in the brainstem and found no significant difference between PLX3397 and vehicle-treated rats (data not shown), which is consistent with other reports (De et al., 2014; Fu et al., 2020; Wyatt-Johnson et al., 2021; Stokes et al., 2022). To further confirm spinal microglial cell depletion, RNA from offspring cervical spinal cord tissue was analyzed for the expression of other genes that are specific to microglia in the CNS **(Fig. 5p)**. Analysis of the integrin alpha M receptor CD11b, the fractalkine receptor CX3CR1 (C-X3-C motif chemokine receptor 1), and the P2Y_12_ purinergic receptor genes by qRT-PCR confirmed significant reductions in their expression in both GNX and GIH offspring (n=5 each, p<0.004 vs. vehicle), indicating a significant loss of microglial cells following PLX treatment.

**Figure 5:**
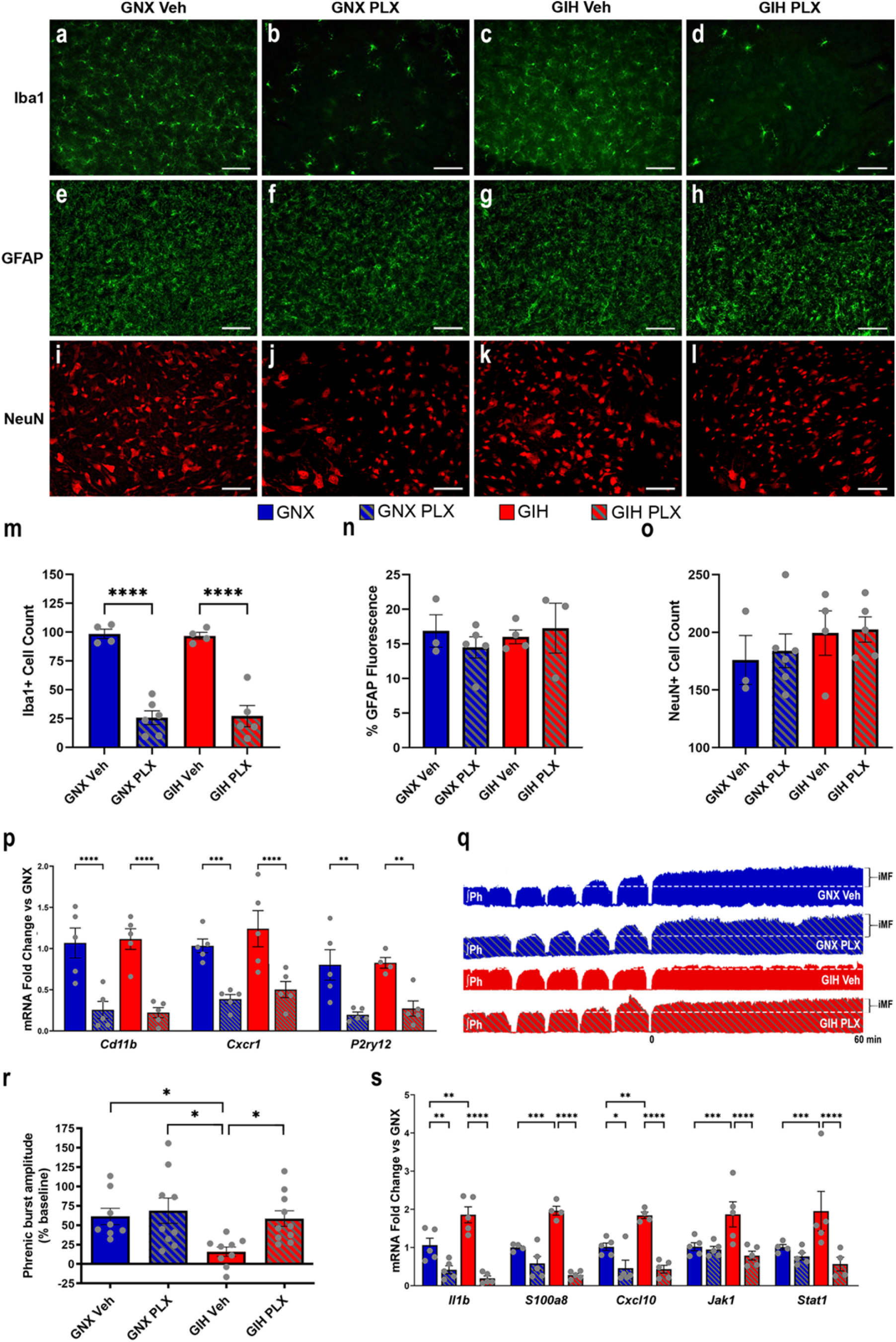
PLX reduces Iba+ cells in cervical spinal cord and restores iMF. **(a-l)** Cervical spinal cord sections from PLX-treated GNX and GIH male rats were immunostained for Iba-1 (microglia) **(a-d)**, GFAP (astrocytes) **(e-h)**, and NeuN (neurons) **(i-l)**. Scale bar = 100um. **(m-o)** Cell count and percent-area fluorescence analyses indicated significant (∼73%) depletion of Iba+ cells in both GNX (p<0.0001) and GIH (p<0.0001) offspring. No significant differences were detected in neurons (GNX p=0.754, GIH p=0.895) or astrocytes (GNX p=0.424, GIH p=0.682). **(p)** Two-way ANOVA revealed PLX treatment resulted in a significant decrease in expression of the microglia-specific genes *Cd11b* (n=5 each; F_(1,16)_=45.94, p<0.0001), *Cx3cr1* (n=5 each; F_(1,16)_=28.12, p<0.0001) and *P2ry12* (n=5 each, except n=4 GIH PLX; F_(1,15)_=25.8, p=0.0001) in both GNX and GIH offspring, confirming evidence that PLX3397 significantly depletes microglia in rats. **(q)** Representative compressed phrenic neurograms depicting phrenic inspiratory output before, during, and for 60 minutes following exposure to recurrent reductions in respiratory neural activity (5, ∼1 min neural apneas). Dotted white line represents baseline amplitude. **(r)** Average percent change (±SEM) in phrenic inspiratory burst amplitude from baseline at 60 minutes following the fifth central apnea. 2-way ANOVA revealed a significant effect of GNX/GIH status (F_(1,32)_ = 5.88, p=0.0211) and vehicle/PLX treatment (F_(1,32)_ = 4.794, p=0.036). Tukey’s post hoc tests revealed that there was no significant difference in the magnitude of iMF between male GNX rats treated with vehicle or PLX3397 (n=8,9 respectively, p=0.9706). Similar to previous results, iMF magnitude in vehicle-treated adult male GIH rats (n=9) was significantly lower than the response in GNX males (p=0.0473). However, PLX-treatment restored the capacity for male GIH (n=10) offspring to trigger compensatory increases in phrenic inspiratory output following recurrent central apneas (p=0.0489 relative to vehicle-treated GIH; p=0.9182 relative to PLX-treated GNX offspring), indicating that microglial depletion rescues the capacity to trigger iMF in male GIH offspring. **(s)** GIH-induced upregulation of *Il1β, S100a8, Cxcl10, Jak1*, and *Stat1* spinal gene expression was also significantly decreased by PLX treatment, suggesting a microglial source of neuroinflammation in the GIH spinal cord [n=4-5; two-way ANOVA with Tukey’s *post-hoc* test). *p<0.05, **p<0.01, ***p<0.001, ****p<0.0001

To test the hypothesis that depletion of inflammatory microglia rescues iMF in adult male GIH offspring (Experimental Series 3), we treated a separate cohort of animals with PLX3397 (**Fig. 5q, r**). PLX3397 treatment did not alter iMF magnitude in male GNX offspring (vehicle: 61.4±10.6% baseline, n=8; PLX: 68.7±16.3% baseline, n=9; p=0.9706). Consistent with our earlier findings (**Figs. 1 and 4**), iMF was significantly impaired in vehicle-treated adult male GIH offspring (15.9±6.1% baseline, n=9) relative to vehicle-treated GNX offspring (p=0.0473; **Fig. 5q, r**). However, microglial depletion with PLX3397 rescued the capacity of adult male GIH offspring to elicit iMF (58.7±10.1% baseline, n=10, p=0.0489 vs. vehicle-treated GIH offspring) to a level similar to that observed in GNX offspring (p=0.9182), indicating that microglial depletion in adulthood can restore the capacity to express iMF in offspring that were exposed to GIH during fetal development. To determine whether microglia-dependent spinal neuroinflammation is also reversed by PLX treatment **(Fig. 5s)**, we analyzed the same inflammatory genes assessed in Fig 2. As before, we found that the inflammatory gene expression that was significantly enhanced in the GIH male cervical spinal cord homogenate (n=4-5; *Il1β* p=0.001; *s100a8* p=0.0006; *Cxcl10* p=0.002; *Jak 1* p=0.0007; *Stat1* p=0.0003) was significantly reduced by PLX treatment, especially in GIH males (*Il1β* GNX p=0.009, GIH p<0.001; *s100a8* GNX p=0.116, GIH p=<0.0001; *Cxcl10* GNX p=0.02, GIH p<0.0001; *Jak 1* GNX p=0.761, GIH p<0.0001; *Stat1* GNX p=0.377, GIH p<0.0001). Collectively, these data show that microglia are crucial mediators of GIH-induced impairment of iMF in adult males and support the idea that although the microglial reprogramming insult occurred *in utero* months earlier, the neuroplasticity deficit can be rescued in the adult male by depleting persistently inflammatory microglia.

### Regulation of physiological variables

**Supplementary Table 1** lists average age and body weight, and average Pa_CO2_, Pa_O2_, pH, mean arterial pressure (MAP) at baseline and 60 min after treatments. Small, but significant, differences in PO_2_ and MAP from baseline to 60 min after the protocol were noted for a few groups, but these values were within normal physiological limits, not associated with iMF magnitude and were consistent with other reports using this anesthetized rat preparation (Dale-Nagle et al., 2011; Streeter and Baker-Herman, 2014b; Baertsch and Baker-Herman, 2015).

## DISCUSSION

We demonstrate that gestational intermittent hypoxia (GIH), a hallmark of maternal SA during pregnancy, reprograms the male offspring neuroimmune system toward enhanced inflammation in CNS regions important in breathing, an effect that lasts into adulthood. Persistent neuroinflammation in male GIH offspring has lasting consequences on the ability to elicit compensatory neuroplasticity in response to a life-threatening event – reductions in respiratory neural activity. Several NF-κB pathway target genes are upregulated in adult male GIH microglia isolated from spinal regions encompassing phrenic motoneurons, with no changes in brainstem microglia. Depleting inflammatory microglia or directly inhibiting spinal NF-κB and STAT transcription factor activation rescued the capacity for adult male GIH offspring to exhibit respiratory plasticity. Adult female GIH offspring did not exhibit neuroinflammation or impaired respiratory neuroplasticity, yet exhibited nearly 4000 differentially expressed genes in spinal microglia, with only a handful of differentially expressed brainstem microglial genes. Mechanisms confering protection to females from detrimental effects of maternal IH are not understood but may involve a protective gene program that mitigates neuroinflammation, and represents an exciting direction for future study. Thus, we demonstrate for the first time a link between maternal IH, sex- and brain region-specific reprogramming of central immune processes, and detrimental effects on respiratory control in adult offspring. Our data indicate that microglia may be a source of cellular memory for *in utero* experiences, releasing neuromodulatory mediators that influence circuits regulating breathing.

Converging evidence suggests that many neurological disorders result from a complex interplay between genetics and early life experiences, particularly in the womb (Bilbo et al., 2018). Several prenatal factors have been implicated, such as maternal infection, stress, obesity, malnutrition, and environmental toxins. Although seemingly diverse, these factors share one commonality: maternal immune system activation. Without question, IH associated with SA causes chronic inflammation in humans and animal models (McNicholas, 2009; Dempsey et al., 2010; Fung et al., 2012; Khalyfa et al., 2017), which is responsible for many morbidities associated with SA. The scientific community has begun to appreciate that maternal SA during pregnancy is detrimental to the health of the newborn (Brown et al., 2018; Johns et al., 2020); however, whether these detrimental effects extend into adulthood is unclear. Correlative evidence suggests that maternal SA during pregnancy may have life-long consequences on offspring neural function in humans. SA is most prevalent in women that are obese or of advanced age, and SA during pregnancy increases risk for gestational diabetes, hypertension, fetal growth restriction, premature birth, NICU admission and lower APGAR scores (Carnelio et al., 2017). These risk factors for, or complications of, SA during pregnancy increase susceptibility to the development of neural disorders in offspring (Krakowiak et al., 2012; Carnelio et al., 2017). Despite this striking correlation, systematic investigations of a mechanistic link between maternal SA and aberrant neural outcomes in offspring are lacking. Indeed, before such a link can be rigorously investigated in humans, animal models must necessarily provide the justification due to the expensive and time-consuming nature of epidemiological studies.

Given that neuroinflammation undermines plasticity in the hippocampus (Rizzo et al., 2018) and respiratory control system (Hocker et al., 2017), we investigated sex-specific differences in inflammatory gene expression in the brainstem and cervical spinal cord, CNS regions associated with the control of breathing, as a mechanism underlying GIH-induced respiratory control deficits. In male GIH offspring, we found enhanced neuroinflammatory gene expression in spinal cord, but not in brainstem, which extends our previous observations of increased neuroinflammation in GIH neonates (Johnson et al., 2018). A tendency towards the opposite (decreased neuroinflammatory gene expression in brainstem and spinal cord) was observed in GIH females. Mechanisms underlying sex-specific effects of GIH are under investigation, but others report sexual dimorphisms in epigenetic gene alterations following prenatal stressors (Nätt et al., 2017; Lei et al., 2020), including gestational sleep fragmentation (Khalyfa et al., 2015).

Epigenetic alterations of the neural immune system resulting from prenatal insults can have long-lasting neurological impacts on offspring (Bergdolt and Dunaevsky, 2019). Having observed spinal neuroinflammation in male GIH offspring, we performed RNA-seq on spinal microglia isolated from male GNX and GIH offspring. Although the mechanistic experiments described here focused on differentially upregulated genes in male GIH offspring spinal microglia related to inflammatory processes, there were also 23 differentially downregulated genes (Table 1). These downregulated genes included *Flt1*, which encodes vascular endothelial growth factor receptor 1 protein, and *Epas1*, which encodes hypoxia inducible factor 2a (HIF-2a) transcription factor. These observations suggest in addition to exaggerated inflammation in male GIH spinal microglia, aberrant function of hypoxia-responsive signaling pathways might also play a role. Additional studies are required to probe functional contributions of downregulated microglial genes in impaired respiratory neuroplasticity in adult male GIH offspring.

Although only six genes were upregulated in spinal microglia from adult male GIH offspring, of the four upregulated genes with known function (*Scarna2, Klkb1, Fcrl2* and *Iκbkε;* **Table 1**), all play roles in inflammation and/or are implicated in neural dysfunction (Chikaev et al., 2005; Khoddami and Cairns, 2013; Hayama et al., 2016; Yin et al., 2020). Recently, NF-κB was found to be upregulated in the nucleus accumbens of mouse offspring whose mothers were injected with poly I:C mid-gestation (Ketharanathan et al., 2021), setting a precedent for dysregulation of offspring NF-κB in models of maternal immune activation. Analyses of upregulated genes from adult male GIH spinal microglia on the ENCODE database indicated that NF-κB and/or STAT transcription factors bind proximally, which guided our subsequent neurophysiological experiments. Local pharmacologic inhibition of NF-κB and STAT transcription factors in the spinal cord rescued expression of compensatory respiratory plasticity in adult male GIH offspring, indicating that region-specific inflammatory signaling plays a role in GIH-induced iMF impairment. Pharmacologic depletion of microglia also rescued iMF, further implicating altered spinal microglial inflammatory function as the underlying mechanism for impaired compensatory respiratory plasticity.

An important caveat to consider when using PLX drugs to reduce microglia number and reactivity is that PLX3397 inhibits CSF1R in all cells, not exclusively in microglia (Kumari et al., 2018; Han et al., 2020). Nevertheless, several lines of evidence indicate that CSF1R inhibition does not impact astrocyte (Qu et al., 2017) or neuronal function (Green et al., 2020), consistent with our findings that astrocyte and neuronal cell numbers were unaffected by PLX treatment, and that phrenic iMF was not impaired in PLX-treated adult male GNX offspring **(Fig. 5)**. While this latter finding shows that phrenic iMF expression does not require microglia for normal expression of plasticity, inflammatory microglia seem sufficient to abolish phrenic iMF in the context of GIH-induced neuroinflammation.

We model SA in pregnancy by delivering IH to pregnant dams during their sleep phase; however, it is acknowledged that SA in humans induces concurrent pathologic conditions besides IH, such as sleep fragmentation, hypercapnia, excessive sympathetic activation, and drastic swings in intrathoracic pressure. Regardless, chronic IH alone replicates many core consequences of SA in humans (Chopra et al., 2016), allowing us to dissociate other concomitant aspects of SA as causal, and simplifying the experimental pathologic insult. Thus, exposing pregnant animals to IH is a logical starting point for investigation into long-lasting effects of SA on offspring, and mechanisms underlying deficits in respiratory control.

Although our studies are the first to show that GIH differentially reprograms adult male offspring spinal microglia to create deficits in the neural control of breathing in adulthood, other studies have interrogated long-lasting consequences of maternal IH on adult offspring physiology. For example, late GIH exposure epigenetically reprograms adipocytes towards a pro-inflammatory phenotype, resulting in offspring metabolic dysfunction (Khalyfa et al., 2017; Cortese et al., 2021). Other studies have found that GIH offspring exhibit endothelial dysfunction (Badran et al., 2019), increased blood pressure (Chen et al., 2018; Song et al., 2021), altered gut microbiome (Cortese et al., 2021), and blunted hypoxic ventilatory responses (Gozal et al., 2003). Although we are beginning to recognize that GIH can have life-long effects on adult physiology, comparatively little is known regarding adult offspring neural function. We recently reported that GIH has sexually dimorphic effects on offspring neural function, and permanently alters the male offspring brain in regions underlying social motivation and cognition, leading to a constellation of deficits that may have relevancy to autism spectrum disorder (Vanderplow et al., 2022). Our current study extends those findings to include CNS regions involved in breathing. GIH transforms developing microglia in male offspring to a pro-inflammatory phenotype present in adulthood, which is linked to respiratory neural dysfunction that results in an inability to respond to reductions in respiratory neural activity with compensatory enhancements in phrenic inspiratory output (i.e., iMF). Strikingly, IH-induced enhancement of microglial pro-inflammatory activity is region specific, apparent in the spinal cord but not in the brainstem. These findings indicate that maternal SA may have a previously unrecognized, detrimental impact on male offspring neural function that may increase vulnerability to developing disorders of ventilatory control associated with breathing instability later in adulthood. Given the stark rise in prevalence of SA during pregnancy in recent years, future work examining additional consequences of GIH-induced microglial reprogramming on male offspring neural function are warranted.

## MATERIALS AND METHODS

All animal experimental procedures were performed according to the NIH guidelines set forth in the Guide for the Care and Use of Laboratory Animals and were approved by the University of Wisconsin-Madison Institutional Animal Care and Use Committee. Sample sizes for each individual experiment is listed in the results section and figure legends.

### Intermittent hypoxic exposures during pregnancy

Timed pregnant Sprague-Dawley rats (gestational age G9) were obtained from Charles River (RRID:RGD_734476; Wilmington, MA, USA) and housed in AAALAC-accredited facilities with 12 h:12 h light-dark conditions. Pregnant rats were purchased in multiples of 2, and were randomly assigned to GNX or GIH exposure. Food and water were provided ad libitum. Beginning at G10, dams were exposed to intermittent hypoxia (GIH) which consisted of alternating 2-minute hypoxic (45 s down to 10.5% O2) and normoxic (15 s up to 21% O2) episodes for 8 h (9:00 am-5:00 pm) daily for 12 days. The apnea-hypopnea index (AHI) of 15 events/h generated by these parameters parallels the kinetics of moderate SA in children and adults (Lim et al., 2015; Brockmann et al., 2016). The control group (GNX) received alternating episodes of room air (normoxia) with the same time and gas flow parameters as GIH dams. Both groups were housed in standard microisolator cages with custom-made acrylic lids to deliver the gases. Rats were removed from the exposure system prior to their expected delivery date (G22), to prevent direct exposure of the pups to IH. Hereafter, offspring of GNX- and GIH-exposed dams are referred to as “GNX” and “GIH” rats, respectively. To control for potential differences in maternal care due to litter size, all litters were reduced to 8 pups (4 males and 4 females per litter if possible) by postnatal day 3. In all experiments, the investigator was blinded to treatment during the study except when populating the plates for qRT-PCR when equal numbers of control samples and sex needed to be included on each plate.

### *In vivo* electrophysiology preparations

Adult rat offspring (8-16 weeks of age) exposed to GNX or GIH were induced with isoflurane (2.5-5%; balance O_2_ and N_2_), tracheotomized, mechanically ventilated (2-4 mL tidal volume; VentElite Small Animal Ventilator, Harvard Apparatus, Holliston, MA, USA), and bilaterally vagotomized to prevent mechanosensory feedback from ventilation. A tail vein catheter was placed for administration of fluids (1:5 sodium bicarbonate/Lactated Ringers Solution) and I.V. drugs. Anesthesia was gradually converted from isoflurane to urethane (1.4-1.7 g/kg, I.V.), and depth of anesthesia was confirmed by monitoring pressor response. Rats were paralyzed with pancuronium bromide (1.0 mg/kg I.V.). Body temperature was maintained between 36-38°C throughout the duration of the surgery and experimental protocol. Blood pressure was monitored, and blood samples were taken periodically from a femoral artery catheter to monitor blood-gas values using an ABL90 Flex (Radiometer, Brea, CA, USA). The phrenic nerve was cut distally, desheathed and inspiratory activity was measured using a bipolar or suction electrode. Raw signals were recorded and digitized with PowerLab data acquisition system (LabChart 8.0, ADInstruments, Colorado Springs, CO, USA), and compound action potentials were amplified (x10k), band-pass filtered (0.3-10 kHz) and integrated (time constant 50 ms). In a subset of rats, an intrathecal catheter was placed for drug delivery. A C2-C3 laminectomy was performed over the spinal midline and a small hole was cut in the dura. A silicone catheter (2 French; Access Technologies, Skokie, IL, USA) connected to a 30 µL Hamilton syringe containing TPCA-1 or vehicle was inserted into the intrathecal space and advanced caudally to lie on the dorsal surface of spinal segment C4.

### Inactivity-induced inspiratory motor facilitation (iMF) protocol

To facilitate rapid induction of brief cessations in respiratory neural activity, rats were slightly hyperventilated and CO_2_ was added to the inspired gas to maintain end-tidal CO_2_ at ∼45 mmHg. One hour following surgery, the CO_2_ threshold for spontaneous breathing was determined (apneic threshold). Inspired CO_2_ was lowered until phrenic activity ceased (apneic threshold), then raised slowly until phrenic activity resumed (recruitment threshold). Inspired CO_2_ was then maintained 2-3 mmHg above recruitment threshold to establish “baseline” respiratory neural activity. Following at least 20 minutes of stable baseline phrenic activity, two arterial blood samples were collected to establish baseline arterial PCO_2_, PO_2_, and pH values. A series of five intermittent central apneas (∼1 min each) were induced by lowering inspired CO_2_ below the apneic threshold until phrenic inspiratory output ceased, then inspired CO_2_ was rapidly returned to baseline levels, at which point phrenic inspiratory activity resumed. Each central apnea was separated by 5 minutes. Importantly, rats remained mechanically ventilated during central apnea and did not experience hypoxia. Following the five central apneas, phrenic activity was recorded for 60 minutes under baseline conditions. Phrenic burst amplitude at 60 minutes post-apneas was compared to baseline phrenic burst amplitude to determine the magnitude of iMF (measured as percent change from baseline). To ensure blood-gas values were maintained at baseline levels following recurrent central apnea, arterial blood samples were drawn at 5-, 15-, 30- and 60-minute after the protocol. After the 60-minute blood draw, rats were euthanized with a urethane overdose. To be included in the analysis, rats had to meet the following criteria: arterial PO_2_ of >150 mmHg, arterial PCO_2_ maintained within 1.5 mmHg of baseline, and base excess within +/-3 mEq/L.

### TPCA-1 treatment

In electrophysiological experimental Series 2, we delivered the Iκ-kinase inhibitor [5-(p-Fluorophenyl)-2-ureido]thiophene-3-carboxamide (TPCA-1; Sigma-Aldrich CAS 507475-17-4; SigmaAldrich, St. Louis, MO, USA) to examine the role of local inflammation in compensatory phrenic facilitation. TPCA-1 was dissolved in DMSO and diluted with artificial cerebral spinal fluid (in mM: 120 NaCl, 3 KCl, 2 CaCl_2_, 2 MgCl_2_, 23 NaHCO_3_, 10 glucose, bubbled with 95% O_2_-5% CO_2_). Vehicle or TPCA-1 (1.4 µg in 10 µL) was delivered in 2 µL boluses over 2 min via an intrathecal catheter, 30-60 min prior to induced neural inactivity.

### PLX3397 treatment

In electrophysiologial experimental Series 3, we pharmacologically depleted microglia cells in the CNS using PLX3397 (MedKoo Biosciences, Morrisville, NC, USA) to examine the role of microglia in iMF expression. This drug selectively kills microglia by inhibiting the tyrosine kinase of the CSF1 receptor, which is a receptor crucial to microglia survival. Each adult offspring rat received daily dosing of PLX3397 or vehicle for 7 consecutive days (80 mg/kg, P.O.). Drugs were formulated in DMSO, 1% PS80, and 2.5% hydroxycellulose.

### Litters used in electrophysiological studies

Electrophysiological experiments included 1-2 offspring of each sex from a single litter. The distribution was as follows: Experimental Series 1: GNX male (11 rats from 9 litters), GIH male (9 rats from 8 litters), GNX female (10 rats from 7 litters), GIH female (9 rats from 9 litters), time control (13 rats from 9 litters); Experimental Series 2: GNX male vehicle (9 rats from 6 litters), GIH male vehicle (8 rats from 5 litters), GNX male TPCA (6 rats from 4 litters), GIH male TPCA (7 rats from 6 litters); Experimental Series 3: GNX male vehicle (8 rats from 7 litters), GIH male vehicle (9 rats from 8 litters), GNX male PLX (9 rats from 8 litters), GIH male PLX (10 rats from 8 litters).

### Immunohistochemistry (IHC) and imaging

In adult offspring, the expression of astrocytes, microglia, and neurons were quantified using IHC following 7 days of PLX3397 treatment. Male GNX and GIH rats (1 rat from a litter) were transcardially perfused with 4% paraformaldehyde (PFA) in 1X phosphate buffered saline (pH 7.4). Cervical spinal cords were collected and post-fixed in 4% PFA for 24 h before being cryoprotected in 20% sucrose solution (1 day) followed by 30% sucrose solution (3 days). Coronal sections 40 µm thick were cut using a sliding microtome (SM200R, Leica Biosystems, Buffalo Grove, IL, USA) in the C3-C5 spinal regions. Spinal slices (n=2-4 from each rat) were incubated with antibodies to IBA1 (1:1000, anti-rabbit, 019-19741, Wako Chemicals, Richmond, VA, USA), GFAP (1: 250, anti-rabbit, ab5804, EMD MilliporeSigma, Burlington, MA, USA), and NeuN (1: 500, anti-mouse, MAV377, EMD MilliporeSigma, Burlington, MA, USA) to identify the expression of microglia, astrocytes, and neurons, respectively. Secondary antibodies were conjugated to Alexa Fluor fluorescent dyes (Invitrogen, Waltham, MA, USA). Images were obtained with a fluorescence microscope (BZX710 series microscope, Keyence, Itasca, IL, USA). Ventral horn images at 20x magnification were taken bilaterally for each slice. IBA+ and NeuN+ cells were hand counted by two blinded scorers using FIJI cell counter (ImageJ, public domain). GFAP+ cells were quantified by comparing percent area fluorescence via FIJI. The analysis threshold was set at default 30, 255.

### Quantitative RT-PCR

Brainstem (medulla and caudal pons) and cervical spinal cord tissues (C3-C6) were isolated (n=5-8/treatment; 1 male and 1 female per litter) and sonicated in Tri-Reagent (Sigma, St. Louis, MO, USA) and stored at −80°C. Total RNA was isolated with the addition of Glycoblue reagent (Invitrogen, Carlsbad, CA, USA) in accordance with the manufacturers’ protocols. Complementary DNA (cDNA) was synthesized from 1.0 µg of total RNA using MMLV reverse transcriptase and a cocktail of oligo dT and random primers (Promega, Madison, WI, USA). qPCR was performed using PowerSYBR green PCR master mix (Thermo Fisher Scientific, Warrington, UK) on an Applied Biosystems 7500 Fast system (Waltham, MA, USA). The ddCT method was employed to determine relative gene expression with respect to 18s ribosomal RNA in brainstem and cervical spinal cord tissue homogenates. The primer sequences used for qPCR are shown in Supplementary Table 2. Primers were designed to span introns wherever possible (NCBI Primer-BLAST) and were purchased from Integrated DNA Technologies (Coralville, IA, USA).

### CD11b+ cell isolation and RNA sequencing

Adult offspring (n=5/treatment; 1 male and 1 female per litter) were euthanized and perfused with cold PBS to remove circulating immune cells from the vasculature of the CNS. Whole brainstems were dissected between the pontomedullary junction and the obex. Spinal cervical C2–C6 vertebrae were removed, and dorsal and ventral C3–C6 cervical spinal segments were extracted based on identification of the spinal roots. Tissues were dissociated into single cell suspensions using papain enzymatic digestion. CD11b+ cells were immunomagnetically isolated as previously described (Crain and Watters, 2009; Nikodemova and Watters, 2012; Crain et al., 2013; Crain and Watters, 2015). Isolated CD11b+ cells will be hereafter referred to as “microglia.”

Total RNA was extracted from freshly isolated microglia with TriReagent according to the manufacturer’s protocol (Sigma-Aldrich, St. Louis, MO) as we have done before (Crain and Watters, 2009; Nikodemova and Watters, 2012; Crain et al., 2013; Crain and Watters, 2015). Total RNA was submitted to Novogene for library construction and paired-end (PE-150) sequencing with an Illumina NovaSeq.

### Maternal care testing

To assess maternal competency of dams exposed to GIH relative to dams exposed to GNX, pup retrieval tests were performed on GIH and GNX litters on postnatal day 3 or 4. All pups were removed from the home cage and placed in under a heating lamp for 10 min before being returned to the home cage with 2 pups placed in the nest with the other pups scattered throughout the cage. Maternal behaviors were observed for 10 min, such as time (latency) to investigate, time to retrieve first pup, time to retrieve all pups, latency to lick, groom, or sniff pups, and time to crouch, burrow, or group pups. Dams that did not retrieve all pups within the 10-min testing period were given a maximal score of 600 s.

### Statistical analyses

Power analyses were done for each experiment based on previous results. All individual datapoints shown in graphs are biological replicates. Technical replicates were also included in the PCR analyses (all samples were pipeted in duplicate). Technical replicates were also used for immunohostochemistry analyses in which 3 tissue sections per animals were quantified and averaged to a single number for each animal.

For all electrophysiology experiments, phrenic nerve burst amplitude was expressed as a percent change from baseline. Phrenic amplitude was measured just prior to blood samples taken at baseline, 15, 30, and 60 min after induced central apnea during the iMF protocol. In experimental series one (Figure 1), statistical differences between groups were determined using Welch’s ANOVA with Dunnett’s T3 multiple comparisons *post hoc* test. In electrophysiological experiments 2 and 3 (Figures 4 and 5), statistical differences between groups were determined using 2-way ANOVA with Tukey’s *post hoc* test. Groups were considered significantly different when *P* values were <0.05. Outliers were determined using Grubbs’ Test with α=0.05. One outlier was identified by the Grubb’s test in the GIH vehicle group in electrophysiological experimental series 2 (Figure 4). For RNA sequencing, index of the reference Rnor6.0 genome was built using Bowtie version 2.2.3. Reads were aligned using TopHat version 2.0.12. Gene counts were made using HTSeq version 0.6.1. Count files were imported to R and filtered such that only genes with a CPM 0.1 expressed in three samples were retained. Counts were normalized using the trimmed mean of M-values method and analyzed for differential expression using EdgeR (Robinson et al., 2010). Differentially expressed genes were identified as statistically significant if the false discovery rate (FDR) was 5%. Results were uploaded to the National Center for Biotechnology Information Gene Expression Omnibus with reference number **GSE142478** (https://www.ncbi.nlm). Transcriptomic Differences in Microglia 211 at ASPET Journals on July 1, 2021 jpet.aspetjournals.org Downloaded from nih.gov/gds). For all qRT-PCR analysis, experimental groups were compared back to their respective control, GNX, using an unpaired t-test with Welch’s correction or a two-way ANOVA with Tukey’s post-hoc test (indicated in the figure legend). No samples for PCR were excluded unless they were identified as outliers by the Grubb’s test, or as outliers by principal component analyses for the RNA-seq dataset. One outlier was identified in each of the the male and female GIH brainstem groups (Fig. 3a and Suppl. Fig. 2a).

## Data availability

All raw data are available on Dryad (https://doi.org/10.5061/dryad.8sf7m0csq) or the National Center for Biotechnology Information Gene Expression Omnibus with reference number GSE142478 (https://www.ncbi.nlm).

## SUPPLEMENTARY FIGURES AND TABLES

**Supplementary Figure 1:**
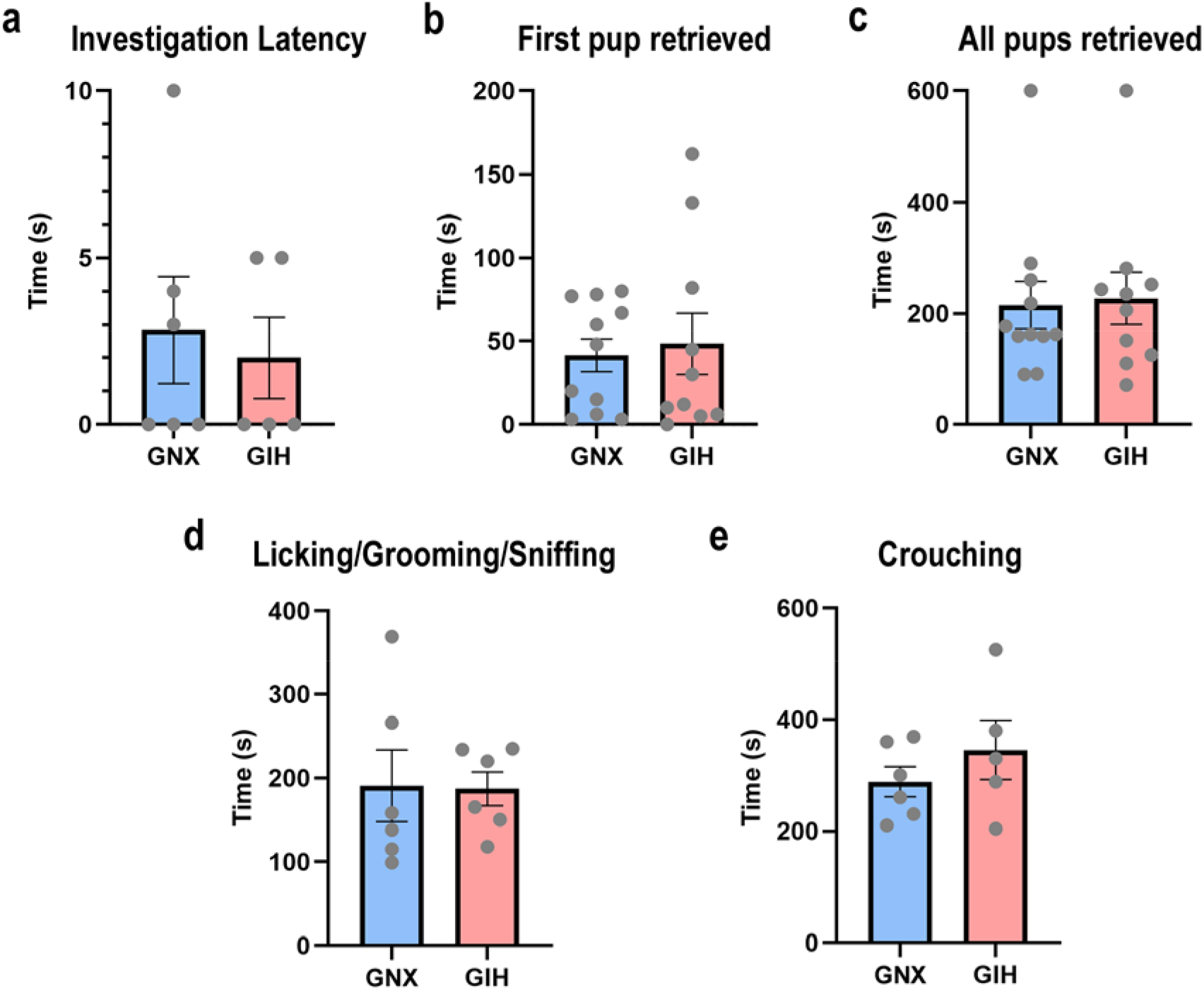
Maternal care is uniform among GNX and GIH dams. Maternal care was assessed in pregnant dams exposed to intermittent hypoxia or normoxia during gestation. No significant differences were observed among several measures used to assess maternal behavior after pup removal and replacement, including the time to investigate pups **(a)**, time to retrieve the first pup **(b)**, time to retrieve all pups into the nest **(c)**, time to lick/groom/sniff pups **(d)**, or time to crouch/nurse pups **(e)**. All p>0.05. Data are displayed as mean ± SEM.

**Supplementary Table 1:** Regulation of physiological variables. Values are expressed as mean ± SEM of arterial pCO_2_, pO_2_, pH, and mean arterial pressure (MAP) at baseline and 60 minutes following reduced respiratory neural activity. Statistics: 2-way repeated measures ANOVA, Holm-Sidak post-hoc comparisons. * Indicates p<0.05 relative to baseline within the treatment.

| Experimental Series 1 |  |  |  |  |  |  |  |  |  |  |
| --- | --- | --- | --- | --- | --- | --- | --- | --- | --- | --- |
|  | GNX males |  | GIH males |  | GNX females |  | GIH females |  | Time controls |  |
|  | baseline | 60 min | baseline | 60 min | baseline | 60 min | baseline | 60 min | baseline | 60 min |
| PaCO<sub>2</sub> (mmHg) | 47.2 ± 1.0 | 47.3 ± 0.8 | 46.2 ± 1.1 | 46.0 ± 1.0 | 51.7 ± 1.0 | 51.9 ± 0.9 | 49.8 ± 1.8 | 49.9 ± 1.9 | 51.2 ± 1.5 | 50.8 ± 1.5 |
| PaO<sub>2</sub> (mmHg) | 243 ± 12 | 236 ± 12 | 257 ± 12 | 245 ± 14 | 249 ± 15 | 201 ± 14* | 231 ± 18 | 207 ± 13* | 245 ± 10 | 225 ± 11* |
| pH | 7.35±0.01 | 7.34±0.01 | 7.35±0.01 | 7.35±0.01 | 7.30±0.01 | 7.29±0.01 | 7.33±0.01 | 7.33±0.02 | 7.31±0.01 | 7.31±0.01 |
| MAP (mmHg) | 133 ± 12 | 115 ± 9 | 145 ± 12 | 134 ± 16 | 145 ± 12 | 134 ± 16 | 101 ± 12 | 91 ± 11 | 150 ± 8 | 137 ± 9 |
| Age (weeks) | 9.1 ± 0.5 |  | 10.6 ± 0.7 |  | 11.9 ± 0.7 |  | 11.3 ± 0.4 |  | 11.0 ± 0.4 |  |
| Weight (g) | 408 ± 19 |  | 477 ± 24 |  | 300 ± 13 |  | 284 ± 11 |  | 363 ± 30 |  |

| Experimental Series 2 |  |  |  |  |  |  |  |  |
| --- | --- | --- | --- | --- | --- | --- | --- | --- |
|  | GNX male vehicle |  | GIH male vehicle |  | GNX male TPCA |  | GIH male TPCA |  |
|  | baseline | 60 min | baseline | 60 min | baseline | 60 min | baseline | 60 min |
| PaCO<sub>2</sub> (mmHg) | 45.8 ± 1.3 | 45.3 ± 1.3 | 45.7 ± 1.1 | 45.2 ± 1.1 | 43.2 ± 0.7 | 43.2 ± 0.9 | 47.2 ± 1.5 | 47.6 ± 1.5 |
| PaO<sub>2</sub> (mmHg) | 252 ± 13 | 233 ± 13 | 263 ± 11 | 250 ± 11 | 261 ± 11 | 246 ± 17 | 285 ± 9 | 275 ± 13 |
| pH | 7.35±0.01 | 7.35±0.01 | 7.35±0.01 | 7.36±0.02 | 7.37±0.01 | 7.38±0.01 | 7.33±0.01 | 7.34±0.01 |
| MAP (mmHg) | 132 ± 11 | 116 ± 10 | 130 ± 8 | 118 ± 7 | 120 ± 8 | 107 ± 8 | 148 ± 6 | 138 ± 7 |
| Age (weeks) | 9.9 ± 0.5 |  | 11.4 ± 0.6 |  | 8.7 ± 0.3 |  | 11.7 ± 1.2 |  |
| Weight (g) | 420 ± 30 |  | 507 ± 31 |  | 382 ± 20 |  | 462 ± 30 |  |

Supplementary Table 1: Regulation of physiological variables
| Experimental Series 3 |  |  |  |  |  |  |  |  |
| --- | --- | --- | --- | --- | --- | --- | --- | --- |
|  | GNX male vehicle |  | GIH male vehicle |  | GNX male PLX |  | GIH male PLX |  |
|  | baseline | 60 min | baseline | 60 min | baseline | 60 min | baseline | 60 min |
| PaCO<sub>2</sub> (mmHg) | 45.3 ± 1.2 | 45.2 ± 1.1 | 45.9 ± 1.4 | 46.3 ± 1.5 | 46.4 ± 1.4 | 46.3 ± 1.3 | 45.4 ± 1.1 | 45.7 ± 1.1 |
| PaO<sub>2</sub> (mmHg) | 258 ± 8 | 243 ± 15 | 233 ± 16 | 236 ± 16 | 272 ± 12 | 235 ± 13* | 263 ± 10 | 255 ± 8 |
| pH | 7.35±0.01 | 7.36±0.02 | 7.35±0.01 | 7.35±0.01 | 7.34±0.01 | 7.35±0.01 | 7.34±0.01 | 7.35±0.02 |
| MAP (mmHg) | 134 ± 8 | 124 ± 10 | 131 ± 9 | 125 ± 7 | 139 ± 9 | 128 ± 9 | 122 ± 8 | 118 ± 8 |
| Age (weeks) | 10.7 ± 0.9 |  | 10.7 ± 0.9 |  | 9.5 ± 0.6 |  | 9.1 ± 0.2 |  |
| Weight (g) | 468 ± 33 |  | 444 ± 31 |  | 409 ± 17 |  | 404 ± 17 |  |

**Supplementary Figure 2:**
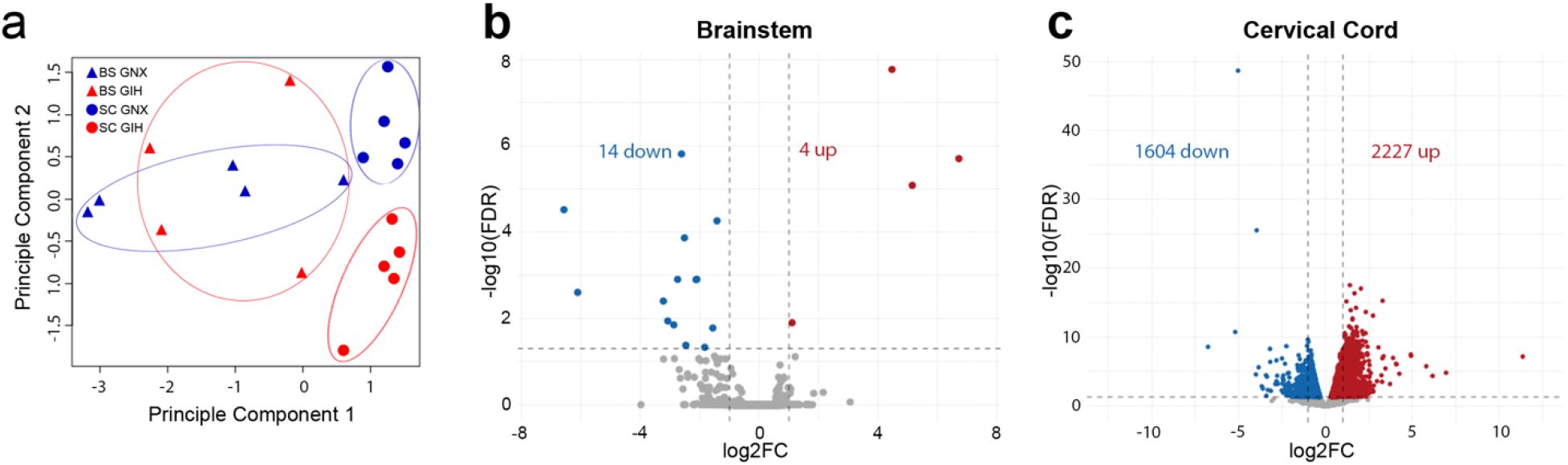
GIH differentially alters the microglial transcriptome in female spinal microglia. **(a)** Principal component analysis of RNA-seq data from female brainstem and cervical spinal microglia (n = 5/treatment) shows transcriptomic differences between brainstem and cervical spinal microglia, and with GIH treatment in spinal microglia. **(b)** Volcano plot indicating differential gene expression in microglia isolated from the female GIH offspring brainstem. In female brainstem microglia, few genes (18) were differentially altered by GIH; red dots represent significantly upregulated genes (FDR < 0.05) and blue dots represent significantly downregulated genes. **(c)** Volcano plot indicating differential gene expression in female cervical spinal microglia from GIH offspring. 3831 genes were differentially expressed; red dots represent significantly upregulated genes (FDR < 0.05) and blue dots represent significantly downregulated genes. FC, fold change.

**Supplementary Table 2:**
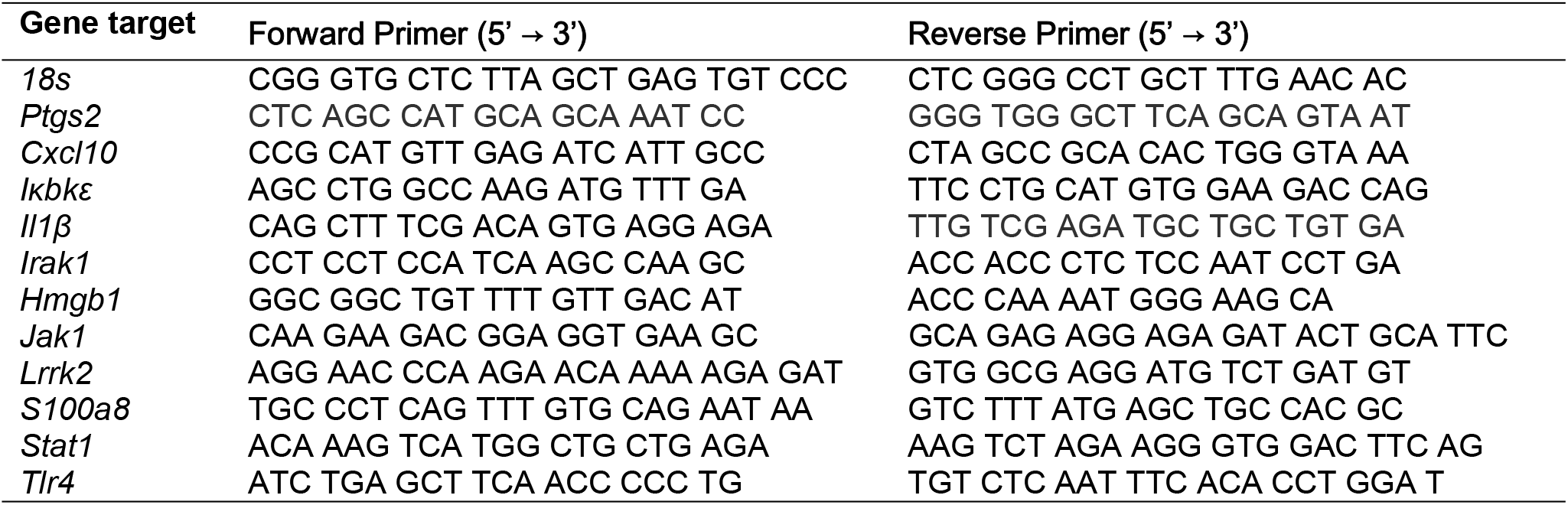
Primer sequences used for quantitative PCR.

## ACKNOWLEDGEMENTS

This work was supported by grants from the National Institutes of Health R01HL105511 (TLB), R01NS085226 (JJW) and R01HL142752 (TLB and JJW). MGG was supported by the National Science Foundation Graduate Research Fellowship Program under Grant No. DGE-1747503. Any opinions, findings, and conclusions or recommendations expressed in this material are those of the author(s) and do not necessarily reflect the views of the National Science Foundation. Support was also provided by the University of Wisconsin-Madison Office of the Vice Chancellor for Research and Graduate Education and Wisconsin Alumni Research Foundation via a UW2020 award (TLB, JJW, MEC, ASR, SMJ).

## AUTHOR CONTRIBUTIONS

CRM, ACE, MGG, ALM, ABR, JNO, BAK conducted experiments and analyzed data. SMJ, ASR, MEC, JJW, TLB planned experiments and analyzed data. SMJ, JJW and TLB contributed to funding acquisition. ASR developed custom bioinformatic tools to analyze RNA-sequencing results. All authors contributed to data interpretation. CRM, ACE, MGG, ALM, ABR, SMJ, JJW, TLB contributed to manuscript preparation. All authors approve the final version of the manuscript.

## REFERENCES

Al-Haddad BJS, Oler E, Armistead B, Elsayed NA, Weinberger DR, Bernier R, Burd I, Kapur R, Jacobsson B, Wang C, Mysorekar I, Rajagopal L, Adams Waldorf KM (2019) The fetal origins of mental illness. Am J Obstet Gynecol 221:549–562.

Almatroodi SA, McDonald CF, Collins AL, Darby IA, Pouniotis DS (2015) Quantitative proteomics of bronchoalveolar lavage fluid in lung adenocarcinoma. Cancer Genomics Proteomics 12:39–48.

Amgalan A, Andescavage N, Limperopoulos C (2021) Prenatal origins of neuropsychiatric diseases. Acta Paediatr 110:1741–1749.

Anderson ME, Buchwald ZS, Ko J, Aurora R, Sanford T (2014) Patients with pediatric obstructive sleep apnea show altered T-cell populations with a dominant TH17 profile. Otolaryngol Head Neck Surg 150:880–886.

Badran M, Yassin BA, Lin DTS, Kobor MS, Ayas N, Laher I (2019) Gestational intermittent hypoxia induces endothelial dysfunction, reduces perivascular adiponectin and causes epigenetic changes in adult male offspring. J Physiol 597:5349–5364.

Baertsch NA, Baker-Herman TL (2015) Intermittent reductions in respiratory neural activity elicit spinal TNF-alpha-independent, atypical PKC-dependent inactivity-induced phrenic motor facilitation. Am J Physiol Regul Integr Comp Physiol 308:R700–707.

Bergdolt L, Dunaevsky A (2019) Brain changes in a maternal immune activation model of neurodevelopmental brain disorders. Prog Neurobiol 175:1–19.

Bilbo SD, Block CL, Bolton JL, Hanamsagar R, Tran PK (2018) Beyond infection - Maternal immune activation by environmental factors, microglial development, and relevance for autism spectrum disorders. Exp Neurol 299:241–251.

Braegelmann KM, Streeter KA, Fields DP, Baker TL (2017) Plasticity in respiratory motor neurons in response to reduced synaptic inputs: A form of homeostatic plasticity in respiratory control? Exp Neurol 287:225–234.

Brockmann PE, Damiani F, Gozal D (2016) Sleep-Disordered Breathing in Adolescents and Younger Adults: A Representative Population-Based Survey in Chile. Chest 149:981–990.

Brown NT, Turner JM, Kumar S (2018) The intrapartum and perinatal risks of sleep-disordered breathing in pregnancy: a systematic review and metaanalysis. Am J Obstet Gynecol 219:147-161.e141.

Carnelio S, Morton A, McIntyre HD (2017) Sleep disordered breathing in pregnancy: the maternal and fetal implications. J Obstet Gynaecol 37:170–178.

Chen L, Zadi ZH, Zhang J, Scharf SM, Pae EK (2018) Intermittent hypoxia in utero damages postnatal growth and cardiovascular function in rats. J Appl Physiol (1985) 124:821–830.

Chikaev NA, Bykova EA, Najakshin AM, Mechetina LV, Volkova OY, Peklo MM, Shevelev AY, Vlasik TN, Roesch A, Vogt T, Taranin AV (2005) Cloning and characterization of the human FCRL2 gene. Genomics 85:264–272.

Chopra S, Polotsky VY, Jun JC (2016) Sleep Apnea Research in Animals. Past, Present, and Future. Am J Respir Cell Mol Biol 54:299–305.

Cortese R, Khalyfa A, Bao R, Gozal D (2021) Gestational sleep apnea perturbations induce metabolic disorders by divergent epigenomic regulation. Epigenomics 13:751–765.

Crain JM, Watters JJ (2009) Cytokine and BDNF expression vary with age and sex in mouse microglia. Journal of Neurochemistry 108:138.

Crain JM, Watters JJ (2015) Microglial P2 Purinergic Receptor and Immunomodulatory Gene Transcripts Vary By Region, Sex, and Age in the Healthy Mouse CNS. Transcr Open Access 3.

Crain JM, Nikodemova M, Watters JJ (2013) Microglia express distinct M1 and M2 phenotypic markers in the postnatal and adult central nervous system in male and female mice. Journal of Neuroscience Research 91:1143–1151.

Dale-Nagle EA, Satriotomo I, Mitchell GS (2011) Spinal vascular endothelial growth factor induces phrenic motor facilitation via extracellular signal-regulated kinase and Akt signaling. J Neurosci 31:7682–7690.

Dalman C, Thomas HV, David AS, Gentz J, Lewis G, Allebeck P (2001) Signs of asphyxia at birth and risk of schizophrenia. Population-based case-control study. Br J Psychiatry 179:403–408.

De I, Nikodemova M, Steffen MD, Sokn E, Maklakova VI, Watters JJ, Collier LS (2014) CSF1 overexpression has pleiotropic effects on microglia in vivo. Glia 62:1955–1967.

Dempsey JA, Veasey SC, Morgan BJ, O’Donnell CP (2010) Pathophysiology of Sleep Apnea. Physiol Rev 90:47–112.

Ding XX, Wu YL, Xu SJ, Zhang SF, Jia XM, Zhu RP, Hao JH, Tao FB (2014) A systematic review and quantitative assessment of sleep-disordered breathing during pregnancy and perinatal outcomes. Sleep Breath 18:703–713.

Fu H, Zhao Y, Hu D, Wang S, Yu T, Zhang L (2020) Depletion of microglia exacerbates injury and impairs function recovery after spinal cord injury in mice. Cell Death Dis 11:528.

Fuller DD, Mitchell GS (2017) Respiratory neuroplasticity - Overview, significance and future directions. Exp Neurol 287:144–152.

Fung AM, Wilson DL, Barnes M, Walker SP (2012) Obstructive sleep apnea and pregnancy: the effect on perinatal outcomes. J Perinatol 32:399–406.

Genest SE, Balon N, Laforest S, Drolet G, Kinkead R (2007) Neonatal maternal separation and enhancement of the hypoxic ventilatory response in rat: the role of GABAergic modulation within the paraventricular nucleus of the hypothalamus. J Physiol 583:299–314.

Gozal D, Reeves SR, Row BW, Neville JJ, Guo SZ, Lipton AJ (2003) Respiratory effects of gestational intermittent hypoxia in the developing rat. Am J Respir Crit Care Med 167:1540–1547.

Green KN, Crapser JD, Hohsfield LA (2020) To Kill a Microglia: A Case for CSF1R Inhibitors. Trends Immunol 41:771–784.

Han J, Fan Y, Zhou K, Zhu K, Blomgren K, Lund H, Zhang XM, Harris RA (2020) Underestimated Peripheral Effects Following Pharmacological and Conditional Genetic Microglial Depletion. Int J Mol Sci 21.

Hayama T, Kamio N, Okabe T, Muromachi K, Matsushima K (2016) Kallikrein Promotes Inflammation in Human Dental Pulp Cells Via Protease-Activated Receptor-1. J Cell Biochem 117:1522–1528.

Hocker AD, Stokes JA, Powell FL, Huxtable AG (2017) The impact of inflammation on respiratory plasticity. Exp Neurol 287:243–253.

Jain SK, Kahlon G, Morehead L, Lieblong B, Stapleton T, Hoeldtke R, Bass PF, Levine SN (2012) The effect of sleep apnea and insomnia on blood levels of leptin, insulin resistance, IP-10, and hydrogen sulfide in type 2 diabetic patients. Metab Syndr Relat Disord 10:331–336.

Johns EC, Denison FC, Reynolds RM (2020) Sleep disordered breathing in pregnancy: A review of the pathophysiology of adverse pregnancy outcomes. Acta Physiol (Oxf) 229:e13458.

Johnson SM, Randhawa KS, Epstein JJ, Gustafson E, Hocker AD, Huxtable AG, Baker TL, Watters JJ (2018) Gestational intermittent hypoxia increases susceptibility to neuroinflammation and alters respiratory motor control in neonatal rats. Respir Physiol Neurobiol 256:128–142.

Ketharanathan T, Pereira A, Reets U, Walker D, Sundram S (2021) Brain changes in NF-κB1 and epidermal growth factor system markers at peri-pubescence in the spiny mouse following maternal immune activation. Psychiatry Res 295:113564.

Khalyfa A, Carreras A, Almendros I, Hakim F, Gozal D (2015) Sex dimorphism in late gestational sleep fragmentation and metabolic dysfunction in offspring mice. Sleep 38:545–557.

Khalyfa A, Cortese R, Qiao Z, Ye H, Bao R, Andrade J, Gozal D (2017) Late gestational intermittent hypoxia induces metabolic and epigenetic changes in male adult offspring mice. J Physiol 595:2551–2568.

Khoddami V, Cairns BR (2013) Identification of direct targets and modified bases of RNA cytosine methyltransferases. Nat Biotechnol 31:458–464.

Krakowiak P, Walker CK, Bremer AA, Baker AS, Ozonoff S, Hansen RL, Hertz-Picciotto I (2012) Maternal metabolic conditions and risk for autism and other neurodevelopmental disorders. Pediatrics 129:e1121–1128.

Kumari A, Silakari O, Singh RK (2018) Recent advances in colony stimulating factor-1 receptor/c-FMS as an emerging target for various therapeutic implications. Biomed Pharmacother 103:662–679.

Lei L, Wu X, Gu H, Ji M, Yang J (2020) Differences in DNA Methylation Reprogramming Underlie the Sexual Dimorphism of Behavioral Disorder Caused by Prenatal Stress in Rats. Front Neurosci 14:573107.

Lim DC, Brady DC, Po P, Chuang LP, Marcondes L, Kim EY, Keenan BT, Guo X, Maislin G, Galante RJ, Pack AI (2015) Simulating obstructive sleep apnea patients’ oxygenation characteristics into a mouse model of cyclical intermittent hypoxia. J Appl Physiol (1985) 118:544–557.

Lockhart EM, Ben Abdallah A, Tuuli MG, Leighton BL (2015) Obstructive Sleep Apnea in Pregnancy: Assessment of Current Screening Tools. Obstet Gynecol 126:93–102.

McNicholas WT (2009) Obstructive sleep apnea and inflammation. Prog Cardiovasc Dis 51:392–399.

Modabbernia A, Mollon J, Boffetta P, Reichenberg A (2016) Impaired Gas Exchange at Birth and Risk of Intellectual Disability and Autism: A Meta-analysis. J Autism Dev Disord 46:1847–1859.

Moore CL, Power KL (1986) Prenatal stress affects mother-infant interaction in Norway rats. Dev Psychobiol 19:235–245.

Nikodemova M, Watters JJ (2012) Efficient isolation of live microglia with preserved phenotypes from adult mouse brain. J Neuroinflammation 9:147.

Nätt D, Barchiesi R, Murad J, Feng J, Nestler EJ, Champagne FA, Thorsell A (2017) Perinatal Malnutrition Leads to Sexually Dimorphic Behavioral Responses with Associated Epigenetic Changes in the Mouse Brain. Sci Rep 7:11082.

Oh SJ, Ahn H, Jung KH, Han SJ, Nam KR, Kang KJ, Park JA, Lee KC, Lee YJ, Choi JY (2020) Evaluation of the Neuroprotective Effect of Microglial Depletion by CSF-1R Inhibition in a Parkinson’s Animal Model. Mol Imaging Biol 22:1031–1042.

Pien GW, Pack AI, Jackson N, Maislin G, Macones GA, Schwab RJ (2014) Risk factors for sleep-disordered breathing in pregnancy. Thorax 69:371–377.

Qu W, Johnson A, Kim JH, Lukowicz A, Svedberg D, Cvetanovic M (2017) Inhibition of colony-stimulating factor 1 receptor early in disease ameliorates motor deficits in SCA1 mice. J Neuroinflammation 14:107.

Rizzo FR, Musella A, De Vito F, Fresegna D, Bullitta S, Vanni V, Guadalupi L, Stampanoni Bassi M, Buttari F, Mandolesi G, Centonze D, Gentile A (2018) Tumor Necrosis Factor and Interleukin-1. Neural Plast 2018:8430123.

Robinson MD, McCarthy DJ, Smyth GK (2010) edgeR: a Bioconductor package for differential expression analysis of digital gene expression data. Bioinformatics 26:139–140.

Song R, Mishra JS, Dangudubiyyam SV, Antony KM, Baker TL, Watters JJ, Kumar S (2021) Gestational Intermittent Hypoxia Induces Sex-Specific Impairment in Endothelial Mechanisms and Sex Steroid Hormone Levels in Male Rat Offspring. Reprod Sci.

Stokes JC, Bornstein RL, James K, Park KY, Spencer KA, Vo K, Snell JC, Johnson BM, Morgan PG, Sedensky MM, Baertsch NA, Johnson SC (2022) Leukocytes mediate disease pathogenesis in the Ndufs4(KO) mouse model of Leigh syndrome. JCI Insight 7.

Streeter KA, Baker-Herman TL (2014a) Decreased spinal synaptic inputs to phrenic motor neurons elicit localized inactivity-induced phrenic motor facilitation. Exp Neurol 256:46–56.

Streeter KA, Baker-Herman TL (2014b) Spinal NMDA receptor activation constrains inactivity-induced phrenic motor facilitation in Charles River Sprague-Dawley rats. J Appl Physiol (1985) 117:682–693.

Vanderplow AM, Kermath BA, Bernhardt CR, Gums KT, Seablom EN, Radcliff AB, Ewald AC, Jones MV, Baker TL, Watters JJ, Cahill ME (2022) A feature of maternal sleep apnea during gestation causes autism-relevant neuronal and behavioral phenotypes in offspring. PLoS Biol 20:e3001502.

Wyatt-Johnson SK, Sommer AL, Shim KY, Brewster AL (2021) Suppression of Microgliosis With the Colony-Stimulating Factor 1 Receptor Inhibitor PLX3397 Does Not Attenuate Memory Defects During Epileptogenesis in the Rat. Front Neurol 12:651096.

Yegla B, Boles J, Kumar A, Foster TC (2021) Partial microglial depletion is associated with impaired hippocampal synaptic and cognitive function in young and aged rats. Glia 69:1494–1514.

Yin M, Wang X, Lu J (2020) Advances in IKBKE as a potential target for cancer therapy. Cancer Med 9:247–258.

